# Alterations in glucocorticoid homeostasis following sleeve gastrectomy

**DOI:** 10.1101/2025.09.01.673462

**Authors:** Seraina O. Moser, Andrei Moscalu, Cullen F. Roberts, Denise V. Winter, Friedrich L. Joos, Cristina Gómez, Eric G. Sheu, Alex Odermatt

## Abstract

Elevated intra-tissue levels of active glucocorticoids in obese patients are associated with insulin resistance, diabetes, impaired immune regulation, and other adverse effects. Sleeve gastrectomy (SG) induces significant weight loss and improves metabolic outcomes, including insulin resistance and hyperlipidemia. Among the enzymes controlling intracellular concentrations of active glucocorticoids, 11β-hydroxysteroid dehydrogenase type 1 (HSD11B1, SDR26C1) has been extensively studied concerning obesity and post-gastric surgery interventions. However, most studies focused on either a single tissue, circulating glucocorticoid levels, or HSD11B1 activity. In this work, we showed that circulating active/inactive glucocorticoid ratio (corticosterone (B)/11-dehydrocorticosterone (A)) negatively correlated with glucose tolerance; SG in male C57BL/6 mice slightly reduced circulating mineralocorticoids and reversed the elevated ratio of B/A seen in sham-operated mice after high-fat diet, while improving glucose tolerance. This change was likely due to increased renal 11β-hydroxysteroid dehydrogenase type 2 (HSD11B2, SDR9C3) activity after SG. To provide a comprehensive overview of the systemic effects post-surgery, we evaluated enzymes involved in glucocorticoid homeostasis across tissues on mRNA, protein, and activity level. Moreover, mRNA expression of transcription factors CCAAT enhancer binding protein alpha (*Cebpa*), *Cebpb* and nuclear factor kappa b (*Nfκb (p50)*), as well as cytokines influencing *Hsd11b1* and *Hsd11b2* gene expression were analyzed. The results emphasize stronger influence of renal HSD11B2 than hepatic HSD11B1 activity on the circulating B/A ratio, supported by intrarenal and intrahepatic B/A ratios. The improved glucocorticoid homeostasis following SG, indicated by decreased B/A ratios, proposes a lower risk for glucocorticoid-mediated adverse health effects, including chronic kidney disease, hypertension, and metabolic disturbances.

**NEW & NOTEWORTHY:** This study revealed improved glucocorticoid homeostasis after sleeve gastrectomy (SG), with lower circulating ratios of active (corticosterone, B)/inactive (11-dehydrocorticosterone, A) glucocorticoids and a better glucose tolerance compared to Sham mice. SG enhanced renal HSD11B2 activity, while hepatic HSD11B1 oxoreductase activity was unchanged. These findings suggest that SG reverses some adverse metabolic effects of high-fat diet by enhancing renal HSD11B2 activity, potentially lowering the risk of chronic kidney disease, hypertension, and metabolic disease.

## INTRODUCTION

### Glucocorticoid synthesis, peripheral and hepato-renal metabolism

Glucocorticoids play crucial roles in various physiological processes, including immune response, stress, and metabolism (1, 2). Increased circulating glucocorticoid levels lead to feedback inhibition of the hypothalamic-pituitary-adrenal (HPA) axis with reduced release of corticotrophin-releasing hormone (CRH) and adrenocorticotropic hormone (ACTH), resulting in decreased glucocorticoid production (3). Several enzymes are involved in adrenal steroidogenesis (**Figure 1**) (4). In humans, cortisol is the main glucocorticoid, with about ten times lower circulating corticosterone levels, whilst rodents lack Cytochrome P450, Family 17, Subfamily A, Member 1 (CYP17A1) expression in the adrenals and produce corticosterone as their main glucocorticoid (4). The adrenals tightly regulate corticosteroid synthesis to maintain circulating concentrations.

**Figure 1:**
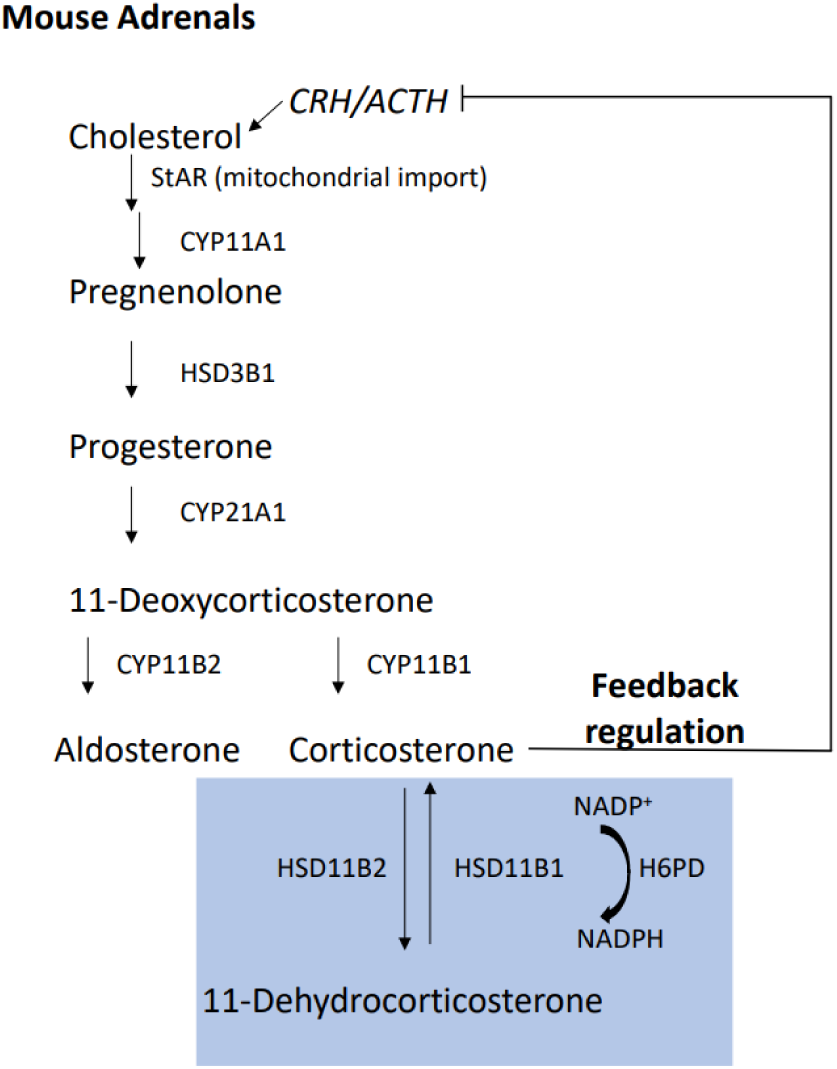
Overview of enzymes involved in adrenal steroid synthesis and peripheral glucocorticoid metabolism in mice. HSD3B1 and CYP21A1 are the mouse homologs of HSD3B2 and CYP21A2 in humans, respectively. CRH = corticotrophin-releasing hormone, ACTH = adrenocorticotropic hormone. Adapted from Aragao-Santiago *et al.* (*105*).

In peripheral tissues, 11β-hydroxysteroid dehydrogenase type 1 (HSD11B1, SDR26C1) and 11β-hydroxysteroid dehydrogenase type 2 (HSD11B2, SDR9C3) regulate the ratio of active (corticosterone (B) in rodents, cortisol (F) in humans) to inactive glucocorticoids (11-dehydrocorticosterone (A) in rodents, cortisone (E) in humans). The multi-functional enzyme HSD11B1 is an endoplasmic reticulum (ER) membrane protein facing the ER lumen, and it acts mainly as an oxoreductase to activate glucocorticoids by using NADPH as cofactor that is supplied by hexose-6-phosphate dehydrogenase (H6PD) (5–8). HSD11B2 is also anchored in the ER membrane, facing the cytosol and catalyzing the reverse reaction to inactivate glucocorticoids in the presence of NAD^+^ (9). HSD11B1 and HSD11B2 control intracellular and intra-tissue concentrations of active glucocorticoids (6).

Glucocorticoids are catabolized in the liver by steroid-5α-reductase (SRD5A1) and steroid-5β-reductase (SRD5B1, aldo-keto-reductase 1D1, AKR1D1) to 5α– and 5β-reduced steroids, representing the first steps of excretion of these glucocorticoid metabolites (10, 11). Additionally, Cytochrome P450, Family 3, Subfamily A, Member 11 (CYP3A11, the mouse homolog of human CYP3A4) catabolizes glucocorticoids to 6β-hydroxysteroids in the liver (12). The metabolized steroids are then glucuronidated or sulfated, followed by excretion via urine and feces. Catabolism affects total circulating corticosteroid levels. **Figure 2** shows the relevant enzymes and steroids analyzed in various tissues of the present study.

**Figure 2:**
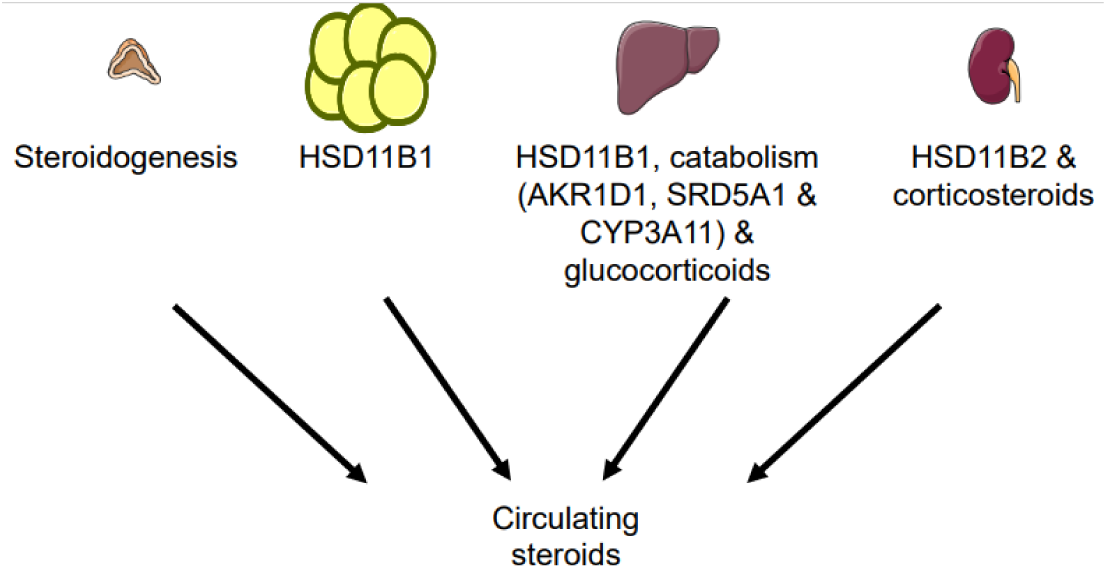
Overview of tissues, enzymes and steroids analyzed in this study. Steroids were measured in serum samples, and to interpret these findings, they were also assessed in several tissues. Steroid synthesis was analyzed in adrenal glands (enzyme expression and corticosteroid levels). Reactivation of glucocorticoids by HSD11B1 was evaluated in epididymal white adipose tissue and in liver. Glucocorticoids and enzymes involved in their catabolism were investigated in liver tissue. In addition, the glucocorticoid-inactivating enzyme, HSD11B2, and corticosteroids were measured in the kidney. Partly created using Servier Medical Art (Servier; https://smart.servier.com/), licensed under a Creative Commons Attribution 4.0 Unported License).

### Glucocorticoids in Obesity

Obesity is a growing, major public health issue worldwide, recognized as a multifactorial disease with a wide array of comorbidities, including type 2 diabetes mellitus (T2DM), hyperlipidemia, hypertension, obstructive sleep apnea, cardiovascular, and chronic kidney disease (CKD) (13, 14). The HPA axis seems to be overactivated in obese individuals and metabolic syndrome patients, with elevated cortisol production that promotes metabolic dysfunction (15, 16). Glucocorticoid excess has been associated with insulin resistance and hyperglycemia, as well as visceral fat accumulation (17–19) with increased circulating free fatty acids and lipid accumulation in skeletal muscle and liver (20–23). Fat cell-conditioned medium promoted the secretion of dehydroepiandrosterone, cortisol, and aldosterone in NCI-H295R adrenocortical cells, implicating an activation of adrenal steroidogenesis that leads to higher cortisol and aldosterone levels in obesity (24).

Besides steroidogenesis, the control of the intracellular concentration of active glucocorticoids in various tissues plays an important role in metabolic diseases. Transgenic mice overexpressing HSD11B1 in adipose tissue develop visceral obesity and metabolic impairments such as insulin-resistant diabetes and hyperlipidemia (25), whereas liver-specific HSD11B1 overexpression led to insulin resistance and hypertension but without obesity (26). In obese Zucker rats, HSD11B1 activity, measured in liver tissue microsomes as the corticosterone to 11-dehydrocorticosterone conversion, was decreased (probably as an adaptive protective mechanism), with no changes observed in subcutaneous adipose tissue (SAT) microsomal preparations, but an increase in omental adipose tissue microsomes compared to the lean group (27). Because of the use of microsomes, this assay is a suitable measure of total actively expressed HSD11B1 protein. In obese patients, elevated *HSD11B1* and *H6PD* mRNA levels were found in adipose tissue (28). Similarly, increased *HSD11B1* mRNA expression was observed in SAT of morbidly obese individuals (29), and HSD11B1 oxoreductase activity measured in SAT biopsies from obese patients was increased and correlated with both overall body fat mass and body mass index (30, 31). This indicates an enhanced glucocorticoid activation in adipose tissue in obesity. Interestingly, in different mouse strains fed a high-fat diet (HFD), a decreased HSD11B1 activity was observed in adipose tissue, suggesting an adaptive protective effect (32) that may be lost in a chronic, low-grade inflammatory, obesity situation.

In livers of morbidly obese patients, *HSD11B1* mRNA levels were positively associated with those of *5α-* and *5β-reductases*, potentially accounting for the lack of systemic hypercortisolism observed in central obesity. Notably, hepatic *5α-reductase* mRNA expression correlated positively with serum insulin levels, which is suggested to enhance hepatic insulin sensitivity (33). Furthermore, hepatic fat content correlated with the urinary ratios of 5β-tetrahydrocortisol (5β-THF)/F and 5β-tetrahydrocortisone (5β-THE)/E, which serve as indicators of 5β-reductase activity (glucocorticoid catabolism) and are linked to elevated fasting plasma insulin levels (30).

The contribution of renal HSD11B2 to systemic glucocorticoid levels seems less clear. Comparisons between severely obese and lean individuals revealed a decreased urinary free F/E ratio, suggesting enhanced HSD11B2 activity, along with a correlation between the urinary F/E ratio and insulin sensitivity (34). Conversely, an analysis of kidney samples from obese and lean Zucker rats, suggested a lower HSD11B2 activity in obese compared to lean rats (27). Collectively, these data indicate a complex regulation of both intra-tissue and systemic concentrations of active glucocorticoids, involving the liver, adipose tissue, and the kidneys. Moreover, caution is warranted when interpreting enzyme activity based on serum and urinary glucocorticoid ratios, as these ratios are influenced by the activities of multiple enzymes (biosynthesis, peripheral and hepatic metabolism, feedback response sensitivity).

### Glucocorticoid changes after bariatric surgery

Bariatric surgery has emerged as a highly effective treatment option for severe obesity and related metabolic disorders. The commonly applied procedures, Roux-en-Y gastric bypass (RYGB), sleeve gastrectomy (SG), and one-anastomosis gastric bypass (OAGB) (reviewed by Salas-Parra *et al.* (35)), result in long-term weight loss and metabolic benefits (36–39).

Shortly after RYGB surgery, patients were reported to have an increase in ACTH levels, with slightly elevated corticosterone levels that did not reach statistical significance. This increase in ACTH suggests an activation of the HPA axis as a stress response to the surgery with enhanced glucocorticoid secretion that normalized thereafter (40). Similarly, serum cortisol and cortisone levels were found to increase compared to pre-operative values in patients undergoing RYGB (15). In morbidly obese individuals, serum cortisol remained elevated up to 6 months after SG (41). Plasma aldosterone concentrations progressively declined from 3 to 12 months following RYGB surgery (42). In HFD-fed mice, SG was shown to normalize the rhythm of circulating glucocorticoid levels compared to sham-operated mice, with corticosterone levels found to be lower than in sham-operated mice in the morning and tending to be higher in the afternoon in SG mice relative to sham-operated controls (43), suggesting an improved glucocorticoid sensitivity and feedback response. Plasma glucocorticoid and aldosterone levels were observed to decrease following OAGB in obese mice, suggesting an amelioration of the overactivation of the HPA axis, with the effect being more pronounced when a longer biliopancreatic limb was applied. A decrease in the plasma B/A ratio further suggested increased HSD11B2 activity (44).

The F/E ratio in SAT biopsies was measured by Woods *et al*. and showed a decrease after RYGB, suggesting lower HSD11B1 activity (15). In female obese patients, *HSD11B1* mRNA levels were found to be elevated in adipose tissue compared to the control group and decreased in a time-dependent manner following surgery (45). Other studies supported the finding of enhanced *HSD11B1* mRNA in SAT in obese patients and a decrease after RYGB (15, 46–48). In VAT, *HSD11B1* mRNA levels were found to be higher in normal-weight patients compared to obese patients, whereas in SAT, similar levels were observed between the groups, with an increase following surgery (49). A decrease in the F/E ratio, with lower *HSD11B1* mRNA levels, was observed in adipose tissue after bariatric surgery in obese patients (48). *HSD11B2* mRNA expression was increased in adipose tissue after fat loss and tended to be lower in non-obese controls compared to post-fat loss patients (48). In SAT and VAT, *HSD11B2* mRNA expression was higher in normal weight individuals compared to obese patients and increased after surgical intervention in SAT (49). This suggests that gastric surgery may lower the concentrations of active glucocorticoids in adipose tissue. Interestingly, prior to gastric bypass, urinary glucocorticoids, including 5α-THF, were found to correlate with waist circumference, suggesting a higher total glucocorticoid biosynthesis with increased hepatic 5α-reductase activity in obesity, which normalized after gastric bypass (47).

Because most of the studies mentioned above focused on either one specific tissue, enzyme or on circulating glucocorticoids, the present study in mice aimed to include several tissues involved in glucocorticoid homeostasis. Additionally, H6PD expression in tissues containing HSD11B1 was examined to provide further insights into the enzyme’s oxoreductase activity. To complement the analysis, CYP3A11, 5α– and 5β-reductases involved in hepatic corticosteroid metabolism, and steroidogenic enzymes expressed in the adrenal glands were also investigated. Besides, corticosteroid hormones were quantified in blood samples as well as liver and kidney tissue samples to obtain information on active and inactive glucocorticoid levels. Together, the results highlight the role of renal HSD11B2 activity for the beneficial effects following SG in mice on HFD.

## MATERIALS AND METHODS

### Animals and diet

Eleven-week-old C57BL6/J male mice were purchased from Jackson Laboratory (strain #000664, Bar Harbor, ME). One cohort was fed a standard chow diet (5% calories from fat; Pico5053; Laboratory Diet, St. Louis, MO) from weaning until 7 weeks of age, when they were reared on an HFD (60% calories from fat; D12492; Research Diets, New Brunswick, NJ). Another 11-week-old C57BL6/J male mouse cohort (standard chow, lean control) was reared on standard chow diet from weaning and through the experiment. Two days pre– and 5 days post-operative, all mice were placed on a DietGel® Recovery (ClearH2O, Westbrook, ME). More detailed food composition can be found in **Suppl. Table 5** – **Suppl. Table 7**. All mice were housed in a climate-controlled environment with a 12-h light/dark cycle, 68-75⁰F and 35-65% humidity, and were acclimated for 1 week before any procedures.

### Sleeve gastrectomy (SG), sham surgery, and functional glucose testing

At 12 weeks of age, all mice were subjected to either SG or sham surgery after weight-matching animals on HFD. Preoperatively, all animals were fasted for 12 h. Anesthesia was induced and maintained using a mix of 95% oxygen and isoflurane (Covetrus, Portland, ME), 3-4% for induction, and 1-3% for maintenance, respectively. After fur removal and cleansing of the anterior abdominal wall skin with povidone-iodine swab sticks (Dynarex, Montvale, NJ, USA), 2% chlorhexidine and 70% alcohol skin wipes (Gama Healthcare, Hertfordshire, UK), and aided by surgical operating loops with a magnification of 2.5x (Designs For Vision Inc, Bohemia, NY), a 5 mm longitudinal, upper midline laparotomy was performed. In the SG group, the stomach was delivered in the wound and reflected to the right and inferiorly to open the angle of His and expose the short gastric vessels, which were controlled using monopolar electrocautery (Bovie Medical Corporation, Clearwater, FL). A longitudinal gastric tube was created by removing 80% of its glandular and non-glandular portions after applying a GIA 45 mm tan linear stapler reload on an Endo GIA™ ultra universal stapler handle (Medtronic, Dublin, Ireland). The sham-operated animals had only the short gastric vessels transected with monopolar electrocautery. The length of the operation was matched for the sham-operated animals to that of SG. For SG mice and sham-operated mice, the peritoneal cavity was closed in anatomical layers with interrupted 6/0 Vicryl (Ethicon, Cincinnati, OH). Periprocedural analgesia was ensured by the administration of buprenorphine SR 0.6 mg/kg/72 h, SC (ZooPharm, Laramie, WY). Postoperatively, all mice were singly housed to accurately monitor food intake (every 3 days), weight (daily for the first week post-op, followed by weekly measurements thereafter), and behavior.

The oral glucose tolerance test (OGTT) was performed following a 4 h fast started at 09:00 am for all cohorts, using 2 mg/g of body weight of 40% glucose solution (Teknova, Hollister, CA, USA), gavaged via a stainless steel, curved, 20 Gauge, 1.5” length, 2.25 mm ball diameter, reusable feeding needle (Roboz Surgical, Gaithersburg, MD, USA). OGTT was performed at 4-week post-op. All mice were placed in new cages at the start of the fast. Baseline glucose was measured for each animal before oral glucose gavage, followed by measurements at 15, 30, 60, and 120 min after gavage. Serum glucose levels were measured using test strips on a Precision Xtra glucometer (Abbott, Alameda, CA, USA).

The experiment was terminated at 5 weeks post-op, starting at 09:00 am and lasting 90 min. Blood and tissues: adrenals, kidneys, liver, and epididymal white adipose tissue (E-WAT) were collected for further *ex vivo* assays.

All procedures were approved by the Institutional Animal Care and Use Committee (IACUC), and all animals were housed and cared for in accordance with institutional and federal guidelines. Animal care complied with the regulations and ethical standards set by Brigham and Women’s Hospital and Harvard University IACUC, the National Institutes of Health (NIH), the American Association for Laboratory Animal Science (AALAS), the Association for Assessment and Accreditation of Laboratory Animal Care (AAALAC), and followed the ARRIVE guidelines and the principles of ethical animal research.

### Gene expression measurements

To determine mRNA expression, tissues (liver 20 ± 5 mg, left kidneys transversally cut, and adrenals 5 ± 2 mg) were homogenized in RLT lysis buffer containing 40 mM dithiothreitol (DTT) using a Precellys-24 tissue homogenizer for 3 x 30 s, at 6’500 rpm and 4°C with Precellys 1.4 mm zirconium oxide beads (P000927-LYSK0-A, Bertin Technologies, Montigny-le-Bretonneux, France). Samples were centrifuged for 10 min at 4°C at 16’000 g. RNA extraction was performed according to the manufacturer using the QIAgen RNeasy Mini Kit (74106, Applied Biosystems, Venlo, The Netherlands) for all tissues except adrenals, for which the QIAgen RNeasy Micro Kit (74004, Applied Biosystems) was used. Adipose tissue (E-WAT), 100 ± 5 mg) was homogenized in TRI Reagent (T9424, Sigma-Aldrich, St. Louis, MO, USA) for 5 x 30 s, at 6’500 rpm and 4°C, followed by adding 200 µL chloroform and subsequent RNA isolation using the QIAgen RNeasy Mini Kit. RNA was quantified with a NanoDrop One© (Witec AG, Sursee, Switzerland) and reverse-transcribed to cDNA using a Takara PrimeScript RT reagent kit (RR037A, Takara Bio, Inc., Kusatsu, Japan). Gene expression was measured on a QuantStudio5 RT System (Thermo Fisher Scientific, Basel, Switzerland). Oligonucleotide primers for RT-qPCR were purchased from Microsynth (Balgach, Switzerland) and are displayed in **Suppl. Table 1**. RT-qPCR was performed with 10 ng of cDNA as a template and 300 nM of oligonucleotide primers and universal SYBR green Master ROX in a total volume of 50 μL (4913914001, Sigma-Aldrich). Runs were started at 95°C for 10 min, followed by 40 cycles (10 s at 95°C, 30 s at 60°C), and completed with a melting curve (95°C for 10 s, 60°C for 30 s and 95°C for 10 s). Ct-values were normalized to the housekeeping gene *18S* rRNA (2−(Ct *gene of interest*−Ct *18S*)).

To determine protein expression, liver (20 ± 5 mg) and E-WAT (100 ± 5 mg) samples were homogenized in RIPA buffer (R8758, Sigma-Aldrich) containing 1x cOmplete protease inhibitor cocktail (11836153001, Roche, Basel, Switzerland) using the Precellys-24 tissue homogenizer as described for mRNA isolation. The supernatant was collected, the protein concentration measured using Pierce™ BCA Protein Assay Kits (23225, Bio-Rad Laboratories, Hercules, CA, USA), and samples were denaturated by adding 4x Laemmli buffer (5 mM Trizma-HCl, pH 6.8, 10% glycerol (v/v), 0.2% sodium dodecyl sulfate (w/v), 1% bromophenol blue (w/v), and 20% β-mercaptoethanol (v/v) followed by boiling at 95°C for 5 min. Samples were stored at –20°C. Per sample, 10 µg protein was separated by SDS-PAGE (10% gel), except for H6PD, where a 7.5% acrylamide gel was used. PVDF membranes (IPVH00010, Merck KGaA, Darmstadt, Germany) were blocked in 10 % milk TBS-T (1.11 g/L Tween-20, 137 mM NaCl, and 20 mM Trizma Base, pH 7.6) at room temperature. Primary antibodies were incubated in 5% milk in TBS-T overnight at 4°C and secondary antibodies for 1 h in 5% milk in TBS-T at RT (for details see **Suppl. Table 2** – **Suppl. Table 4**). Immobilon Western horseradish peroxidase (HRP) substrate kit (WBKLS0500, Merck KGaA) was used to visualize antibody-labeled proteins in a Fusion FX (Vilber Collégien, France). Fusion FX-associated FusionCapt Advance software was used to assess relative protein expression by densitometry.

### HSD11B1 and HSD11B2 activity measurements

Hepatic HSD11B1 activity was determined as described previously with minor modifications (50). Briefly, liver tissue (55 ± 5 mg) was cut and homogenized in 1 mL buffer (20 mM Trizma base (pH 7.5), 50 mM KCl, 2 mM MgCl_2_, 0.25 M sucrose, 1x cOmplete protease inhibitor cocktail (Roche)) with Precellys 1.4 mm zirconium beads in a Precellys-24 tissue homogenizer (3 x 60 s at 6’500 rpm and 4°C). Samples were centrifuged for 20 min at 4°C at 16’000 g. Supernatants were transferred to 1.5 mL conical tubes and sonicated (HIEL_17006, UP50H, Hielscher Ultrasonics, Teltow, Germany), 10-15 times (0.3 cycle, 20% amplitude). Protein concentration was measured using the Pierce™ BCA protein assay kit (Bio-Rad Laboratories). To measure activity, 1.1 µg/µL liver protein was incubated with 10 nCi tritium-labeled cortisone, 190 nM cortisone, and 500 µM NADPH in homogenization buffer in a thermoshaker for 30 min at 37°C. To stop the reaction, 10 µL of cortisol and cortisone, 2 mM each, in methanol was added. Steroids were separated by TLC and analyzed by scintillation counting to determine HSD11B1 activity as previously described (50).

Renal HSD11B2 activity assay was measured as described earlier with minor modifications (51). Briefly, the right kidney was transversally cut and homogenized (1.4 mm zirconium beads, Precellys-24 tissue homogenizer (3 x 60 s at 6’500 rpm and 4°C)) in buffer (1 mL, 20 mM Trizma-HCl (pH 8), 10 mM HEPES, 0.25 M sucrose, 1x cOmplete protease inhibitor cocktail), followed by centrifugation (20 min, 4°C for 16’000 g). Supernatants were sonicated (10-15 times, 0.3 cycle, 20% amplitude, HIEL_17006. UP50H, Hielscher Ultrasonics). Protein concentration of the samples was analyzed (Pierce™ BCA protein assay kits) and 1.8 µg/µL kidney protein was incubated in a thermos-mixer for 60 min at 37°C, with 10 nCi tritium-labeled cortisol, 40 nM cortisol and 500 µM NAD^+^ in reaction buffer (300 mM NaCl, 1 mM EDTA, 1 mM EGTA, 0.25 M sucrose, 20 mM Trizma-HCl, pH 8.0). Reactions were stopped (10 µL each of 2 mM cortisol and cortisone in methanol), steroids separated by TLC, followed by scintillation counting and assessment of HSD11B2 activity as previously described (51).

### Analytical measurements

Serum concentrations of corticosterone, 11-dehydrocorticosterone, 11-deoxycorticosterone, aldosterone, androstenedione, testosterone, and progesterone were analyzed as described earlier with minor modifications (52). Volume-matched calibrators prepared in charcoal-stripped mouse serum were used for calibration and subsequently treated as samples. For solid-phase extraction (SPE), each sample (100 µL) was mixed with a protein precipitation solution (100 µL, 0.8 M zinc sulfate in methanol/water (50/50, v/v) containing labeled internal standards (ISTD): aldosterone (D7, 66 nM), corticosterone (D8, 66 nM), androstenedione (D7, 23 nM), testosterone (C13-3, 21 nM) and progesterone (D8, 26 nM)). Prior to SPE, all samples were diluted with water to a final volume of 1 mL and incubated (10 min, 4°C, 1’300 rpm). Samples were centrifuged (10 min, 16’000 g, 4°C) and supernatants (950 µL) transferred to preconditioned (methanol, water, 3 mL each) Oasis HBL (3cc) cartridges (Waters, Milford, MA, USA, Distributor: Morvay Analytik). Cartridges were washed with water and 10% methanol in water (1:9, v/v, 3 x 1 mL each). Steroids were eluted with methanol (2 x 0.75 mL), evaporated to dryness (Genevac EZ-2 (SP Scientific, Warminster, PA, USA) and re-constituted in methanol (25 µL, 10 min, 4°C, 1’300 rpm). Analyte separation was achieved using a reverse-phase column (1.7 µm, 2.1 mm x 150 mm; Acquity UPLC BEH C18; Waters, Distributor Morvay Analytik) with an injection volume of 5 µL. Steroid content was analyzed by liquid chromatography coupled with tandem mass spectrometry (LC-MS/MS) using an Agilent 1290 Infinity II UPLC connected to an Agilent 6495 triple quadrupole mass spectrometer equipped with a jet-stream electrospray ionization interface (Agilent Technologies, Santa Clara, CA, USA). Data acquisition and quantitative analyses were performed using MassHunter software (Version B.10.0. Build 10.0.27, Agilent Technologies).

Due to the absence of a matrix surrogate, hepatic corticosterone and 11-dehydrocorticosterone levels were measured against a volume-matched calibration curve prepared in water:0.5% formic acid (v/v, 300 µL). Calibrators were subsequently treated as samples. Liver tissue (30 ± 5 mg) was cut and homogenized in hexane:ethyl acetate (1:1, v/v, 1 mL) containing ISTD in PBS (100 µL corticosterone (D8, 66 nM)) with a Precellys-24 tissue homogenizer (6’500 rpm, 3 x 30 s, 4°C, 1.4 mm zirconium beads). For liquid-liquid-extraction (LLE) samples were diluted with 0.5% formic acid in water (v/v, 400 µL) while shaking (10 min, 25°C), followed by centrifugation (10 min, 18’200 g). LLE was repeated twice and organic fractions (2x 800 µL) were collected and evaporated to dryness at 35°C (Genevac EZ-2, SP Scientific). Samples where resuspended in hexane by sonication (300 µL, 10 min) and steroids LLE extracted from hexane with methanol/water/0.5% formic acid (1:1, v/v, 1 mL) for 10 min and centrifuged (10 min, 18’200 g). Samples were diluted with water:0.5% formic acid to a final volume of 1 mL. SPE was performed on preconditioned (methanol, water, 3 mL each) Oasis HLB 3cc columns. Samples (1’900 µL) were loaded on the cartridges and washed with 0.5% formic acid in water and water (3 x 1 mL each), followed by drying for 3 min.

Steroids were eluted by methanol (2 x 700 µL), samples evaporated to dryness, and resuspended in methanol (50 µL). Steroids were separated and analyzed as described above for serum steroids.

Similarly, renal corticosterone, 11-dehydrocorticosterone, 11-deoxycorticosterone, aldosterone, androstenedione, testosterone, and progesterone were measured against a volume-matched calibration curve prepared in PBS and calibrators were subsequently treated as samples. Left kidney samples, 30 ± 5 mg, were homogenized twice in 800 µL chloroform/isopropanol, 1:1, v/v, containing ISTD solution: aldosterone (D7, 66 nM), corticosterone (D8, 66 nM), androstenedione (D7, 23), testosterone (C13-3, 21 nM) and progesterone (D8, 26 nM), dissolved in protein precipitation solution (100 µL, 0.8 M zinc sulfate in water/methanol; 50/50, v/v). Samples were centrifuged at 16’000 g for 10 min and the combined supernatant (1.6 mL/sample) was evaporated to dryness in a Genevac EZ-2 at 35°C. Prior to SPE, samples were resuspended with PBS (100 µL), vortexed, and sonicated (10 min, 25°C). Samples and calibrators were diluted with water to a final volume of 1 mL, followed by SPE extraction on pre-conditioned (methanol, water, 3 mL each) Oasis HLB 3cc columns. Samples (950 µL) were loaded to columns, washed with 10% methanol in water and water (3 x 1 mL each), and subsequently dried for 8 min. Steroids were eluted in methanol (2 x 750 µL) and evaporated to dryness prior to resuspension in methanol (25 µL). Steroids were separated and analyzed as described above for serum steroids.

Intrahepatic bile acids were quantified as described previously (53). Liver tissues (30 ± 5 mg) were homogenized using Precellys 1.4 mm zirconium beads as described above. Samples were extracted using 1 mL water:chloroform:methanol (1:1:3, v/v/v, supplemented with deuterated standards), incubated for 15 min at 37°C and 850 rpm in a thermoshaker, and centrifuged for 10 min at 20°C and 16’000 g. Supernatants (800 µL) were transferred to new tubes and the extraction process was repeated once more. Samples (1.6 mL) were evaporated to dryness in a Genevac EZ-2 at 35°C and reconstituted in 200 µL methanol:water 1:1 (v/v). Bile acids were separated by an Acquity UPLC BEH C18 reversed-phase column and analyzed using an Agilent 1290 UPLC connected to a 6495 MS/MS. The injection volume was 3 µL.

### Cytokine measurements in kidney tissue samples

Cytokines in kidney tissues were assessed using the LEGENDplex™ Mouse Inflammation Panel (13-plex) (740446, Biolegend, Amsterdam, Netherlands) and measured on a CytoFLEX S Flow Cytometer (Beckman Coulter, Nyon, Switzerland). Data were analyzed using Data Analysis Software Suite for LEGENDplex™ (Biolegend). Kidney samples were prepared as described by Hamid *et al.* (54). Right kidneys were transversally cut and homogenized as described above for steroid analysis in 500 µL buffer (pH 7.4) containing 25 mM Trizma base, 150 mM NaCl, 1 mM EDTA, 1 mM EGTA, 1% IGEPAL, and 1x cOmplete protease inhibitor cocktail (Roche) per 10 mL of buffer. Samples were centrifuged for 15 min at 4°C and 16’000 g, protein concentration determined with Pierce™ BCA protein assay kit (Bio-Rad Laboratories), and 20 µg/µL of kidney homogenate was used for cytokine quantification.

### Statistics

Statistical analysis were performed with GraphPad Prism Software 10.2 (RRID: SCR_002798, San Diego, CA, USA), using one-way ANOVA with Tukey’s multiple comparison test (respective p-values are indicated). Outliers were identified and excluded using Grubbs’ test (α=0.05). Data represent mean ± SEM.

## RESULTS

### Improved glucose tolerance post-SG

Mice that underwent sham surgery (Sham) or sleeve gastrectomy (SG) were put on HFD throughout the experiment. The lean control (LC) group was age-matched to Sham and SG mice, sham-operated, but fed a standard chow diet. Following the surgical procedure, an initial decrease in body weight was observed in both Sham and SG mice, with a more pronounced drop in the latter (**Figure 3** a/b). The percentage of weight change increased in all three groups after eight days, with a comparable weight gain for Sham and SG mice but a less pronounced increase for the LC group (**Figure 3** b). Importantly, the impaired oral glucose tolerance and elevated glucose levels after a challenge seen in the Sham group were reversed in the SG group (**Figure 3** c). The area under the curve (AUC) of the OGTT revealed increased levels for the Sham mice (under HFD) compared to LC (under standard chow) and SG mice (under HFD) (**Figure 3** d) (p < 0.0001).

**Figure 3:**
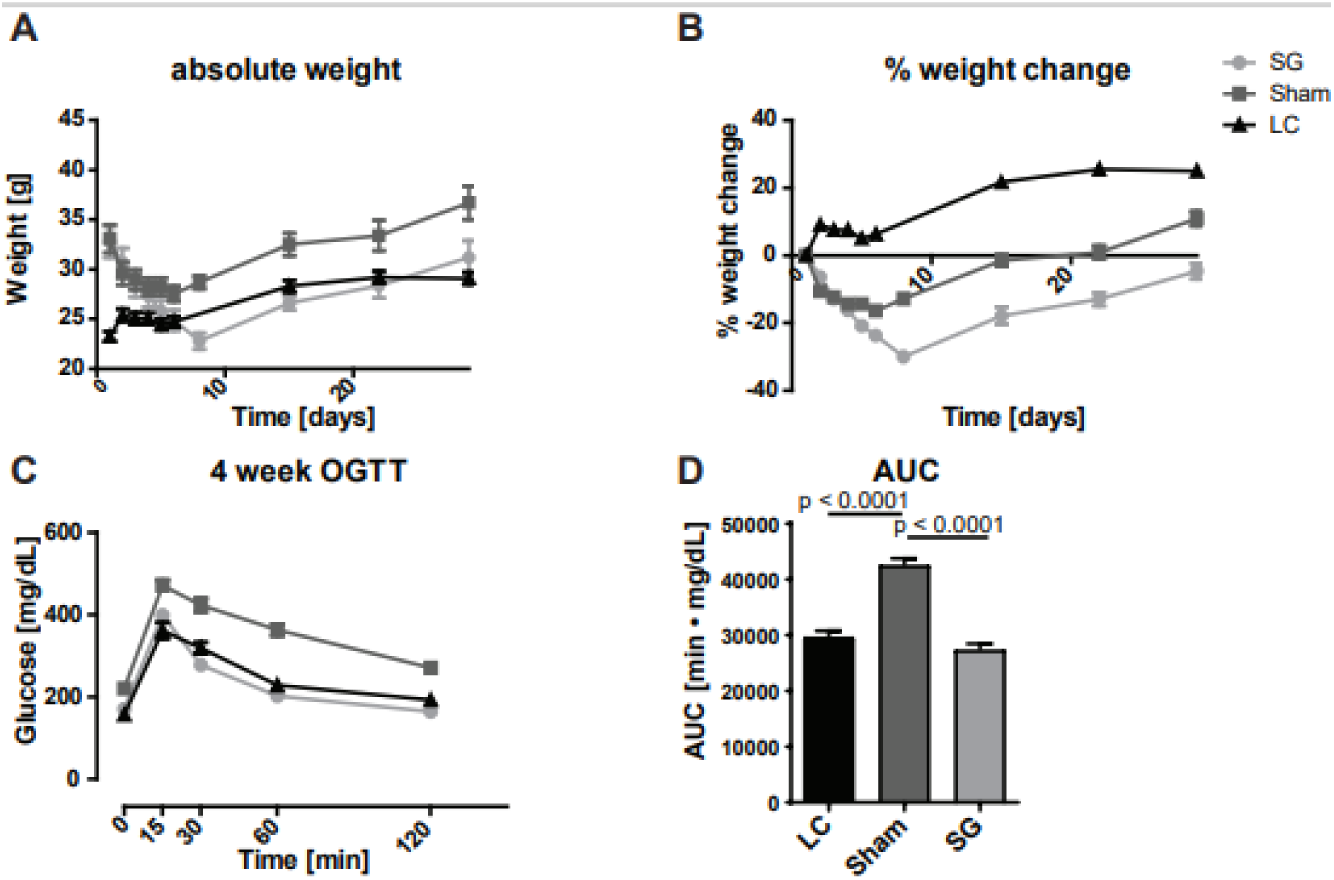
Effects of diet and SG on body weight and oral glucose tolerance. Absolute (a) and percentage weight changes (b) in LC on standard chow, Sham mice on HFD, and mice after SG on HFD. Oral glucose tolerance tests (OGTTs) were performed four weeks after SG (c), and the AUC values of the OGTT of LC, Sham, and SG mice were determined (d). Statistics: One-way ANOVA with Tukey’s multiple comparison test; p-values are indicated. Values represent mean ± SEM. AUC = Area Under the Curve. LC = Lean Control (n=8, standard chow), Sham (n=8, HFD), and SG = Sleeve Gastrectomy (n=10, HFD), HFD = High-fat diet.

### Improved circulating and intra-tissue glucocorticoid levels following SG

Compared to the LC group that was fed a standard chow, sham-operated mice on HFD (Sham) showed approximately 50% lower serum corticosterone (B) concentrations (p = 0.0292), while SG was not altered compared to the LC group was (ns, p = 0.1617) (**Figure 4** a). Circulating 11-dehydrocorticosterone (A) concentrations showed a similar pattern, with a more pronounced decrease in the Sham group to about 20% of the levels seen in LC mice (p < 0.0001). The B/A ratio, a marker for active/inactive glucocorticoids, which is inversely associated with HSD11B2 activity, was elevated in Sham mice and tended to be reversed post-SG (ns, p = 0.0567). Importantly, the serum B/A ratio positively correlated with blood glucose levels at 120 min after OGTT (p = 0.046) and the AUC (p = 0.0269). The mineralocorticoids aldosterone and 11-deoxycorticosterone (11-DOC) were both lower in serum of Sham and SG mice compared to the LC group, but SG had only minor lowering or no effect on aldosterone and 11-DOC, respectively (**Figure 4** a).

**Figure 4:**
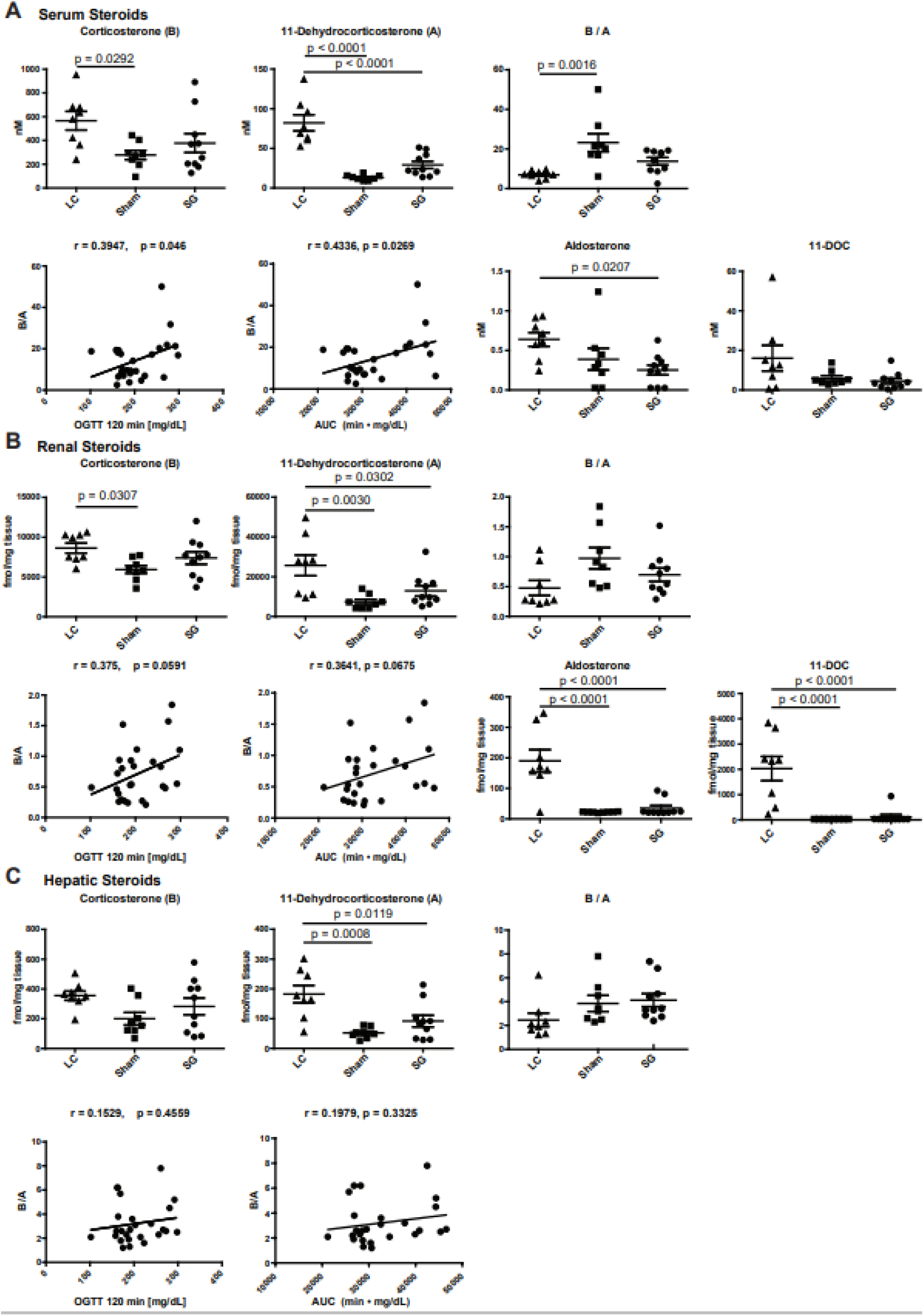
Effects of HFD in Sham and SG mice on corticosteroid metabolism compared to LC mice fed standard chow. Circulating (a) and intrarenal (b), as well as intrahepatic (c) corticosteroids were measured by LC-MS/MS. Linear regression was applied between B/A and OGTT at 120 min, as well as B/A and AUC, and correlation calculated using GraphPad Prism version 10.2. Statistical analysis were performed using one-way ANOVA with Tukey’s multiple comparison test; p-values are indicated. Values are displayed as mean ± SEM. LC = Lean Control (n=8, standard chow), Sham (n=8, HFD), and SG = Sleeve Gastrectomy (n=10, HFD), HFD = High-fat diet, AUC = Area Under the Curve.

To assess the contributions of renal HSD11B2 and hepatic HSD11B1 activities to the concentrations of circulating glucocorticoids, intra-tissue corticosteroid levels were determined. The pattern of intrarenal B and A resembled that observed in serum, with decreased B and A in the Sham compared to the LC group and a trend reversal following SG. The intrarenal B/A ratio increased in Sham mice and showed a tendency to recover after SG. The intrarenal B/A ratio tended to positively correlate with blood glucose levels at 120 min after OGTT (p = 0.0591) and the AUC (p = 0.0675). Interestingly, intrarenal aldosterone and 11-DOC levels dropped dramatically under HFD, whereby SG did not affect this decrease (**Figure 4** b).

The intrahepatic B levels tended to decrease in Sham mice compared to LC mice, with a trend to reverse after SG. Similarly, intrahepatic A levels decreased in Sham compared to LC mice (p = 0.0008), with reversal after SG (p = 0.0119). Despite the lower intrahepatic glucocorticoid levels under HFD, the ratio of active to inactive glucocorticoids was higher. However, SG did not affect the intrahepatic B/A ratio and B/A did not correlate with blood glucose concentration after OGTT (120 min) or the AUC (**Figure 4** c).

No significant changes in serum testosterone, androstenedione, and progesterone concentrations were observed between LC, Sham, and SG groups (**Suppl. Figure 1**), suggesting no changes in gonadal steroid production.

Together, these findings suggest a rebalancing by SG of the impaired glucocorticoid homeostasis caused by HFD, supported by at least a partial normalization of serum and intra-tissue levels of active glucocorticoids post-SG. To begin to understand the observed alterations in corticosteroid levels and B/A ratios, the expression of HSD11B1 and HSD11B2, and related genes were assessed in different tissues.

### Changes in renal HSD11B2 expression following SG

The kidney tissue weight of Sham and SG mice under HFD tended to be slightly higher than that of LC mice (**Figure 5** a). Whilst *Hsd11b2* mRNA expression and HSD11B2 activity were not different between LC and Sham mice, an increase was observed after SG, with significantly higher enzyme activity for SG compared to LC mice (p = 0.0252) (**Figure 5** b). This provides an explanation for the lower B/A ratios seen in serum and kidney tissue for SG compared to Sham mice (**Figure 4** a/b). Interestingly, the transcription factors CCAAT enhancer binding protein beta (*Cebpb*) and nuclear factor kappa b (*Nfκb (p50)*), both reported to induce *Hsd11b2* expression (55, 56), were increased following SG compared to Sham and LC mice (**Figure 5** c). The analysis of proinflammatory cytokines associated with NFκB activation yielded an inconclusive pattern. Whilst HFD increased the mRNA expression of Tumor-necrosis factor alpha (*Tnf-α)* and Interleukin 6 (*Il-6*), with higher levels after SG, it slightly lowered protein levels with a very slight trend increase in SG compared to Sham mice (**Figure 5** d/e). In addition, HFD significantly lowered IL-1β protein levels in Sham and SG compared to LC mice, with a slight trend increase in SG compared to Sham mice; however, IL-1α protein significantly increased after SG compared to both LC and Sham mice (**Figure 5** e/f). Furthermore, other cytokines were measured in this panel, showing a trend decrease in Sham compared to LC mice and a tendency of reversal after SG (**Suppl. Figure 2**).

**Figure 5:**
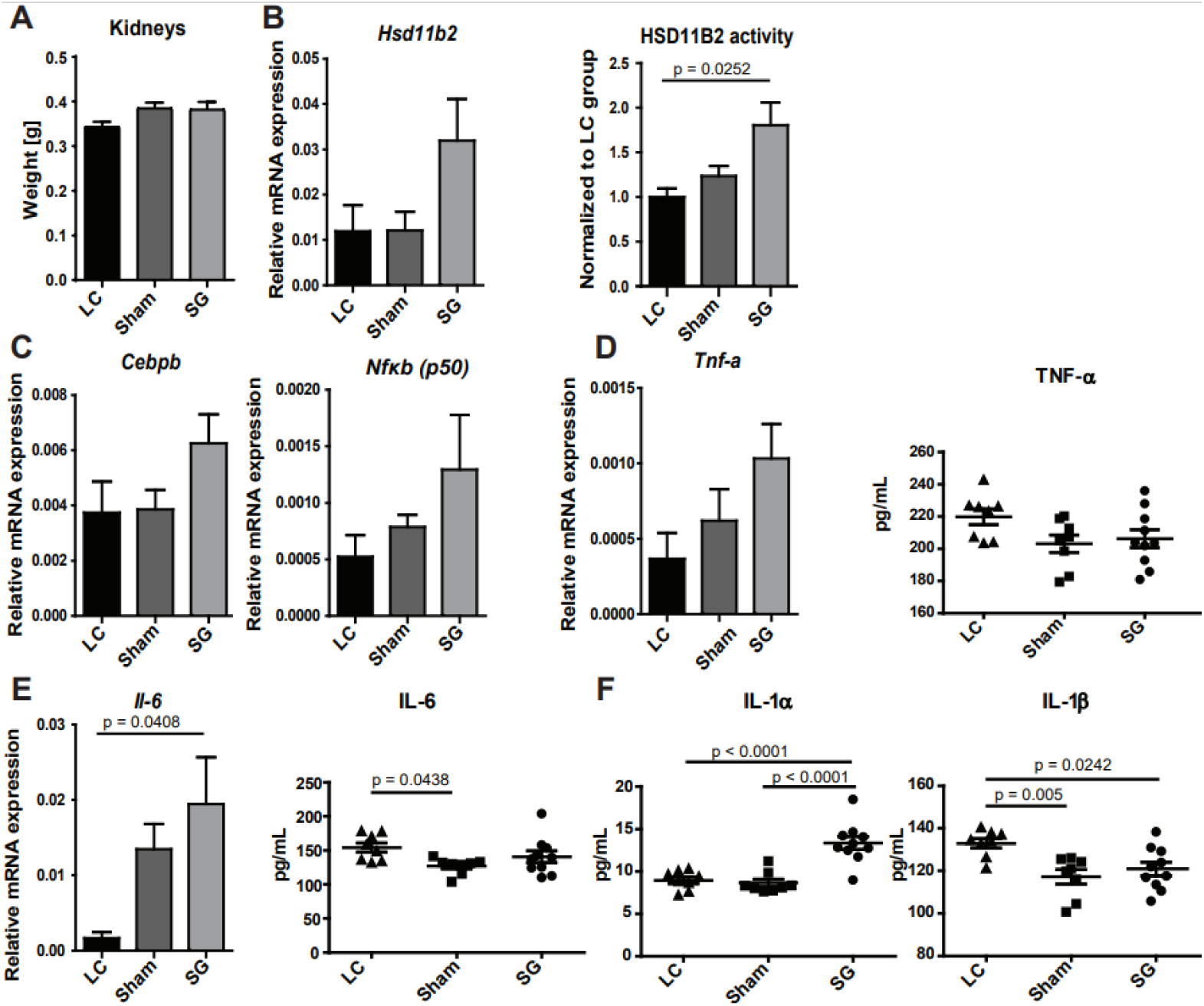
Renal HSD11B2 expression, transcription factors, and proinflammatory cytokines following SG. Kidney tissue weight of LC (fed standard chow) and Sham and SG mice on HFD (a). Measurements of mRNA levels by RT-qPCR, normalized to *18S* rRNA (b-d). Determination of HSD11B2 activity in cell lysates by measuring the conversion of radiolabeled cortisol to cortisone. Values were normalized to the respective LC control group within each reaction. LC1-4 served as controls for Sham183-186 and SG295-297, 299, 301; LC5-8 served as controls for Sham187-190 and SG302-306 (b). Protein levels of cytokines were measured by flow cytometry (LEGENDplex™ Mouse Inflammation Panel (13-plex)) (d-f). Statistical analysis were performed using one-way ANOVA with Tukey’s multiple comparison test; p-values are indicated. Values are displayed as mean ± SEM. LC = Lean Control (n=8, standard chow), Sham (n=8, HFD), and SG = Sleeve Gastrectomy (n=10, HFD), HFD = High-fat diet.

### Alterations of gene expression in epididymal white adipose tissue due to HFD and SG

Excessive accumulation of visceral fat is a co-morbidity of obesity and the metabolic syndrome and it is strongly associated with hypertension (15, 25, 57). The analysis of E-WAT weight revealed a 4 to 5-fold increase in Sham mice under HFD that was significantly lowered post-SG (**Figure 6** a). The mRNA expression of fatty acid synthase *Fasn* and hormone-sensitive lipase *Lipe*, two key genes involved in fatty acid metabolism, showed similar patterns (**Figure 6** a). The mRNA expression of *Hsd11b1*, *H6pd,* and *G6pt* showed similar but less pronounced changes with a trend increase for Sham compared to LC mice, and a partial reversal after SG (**Figure 6** b-d). This was also seen for the *Cebpa*, *Cebpb* and *Nfκb (p50)* mRNA expression, transcription factors known to regulate *Hsd11b1* transcription (**Figure 6** e), and for the proinflammatory cytokine *Tnf-a*, whilst *Il-6* mRNA tended to increase with HFD, and even further after SG (**Figure 6** f). Whilst the H6PD and G6PT protein expression patterns resembled those of the mRNA expression, HSD11B1 expression was almost completely abolished by HFD in the Sham group and partially reversed following SG (**Figure 6** b).

**Figure 6:**
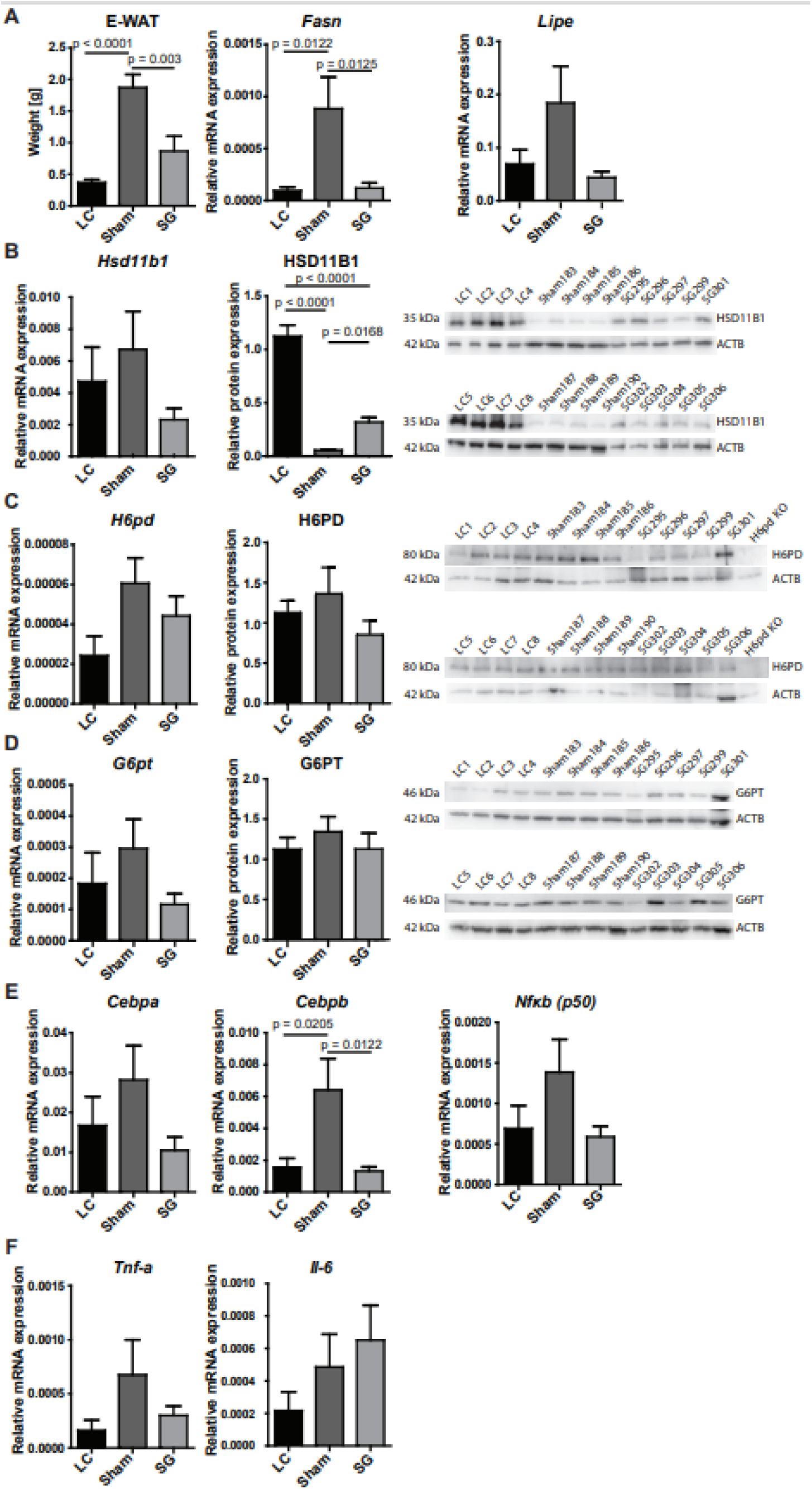
Impact of HFD and SG on E-WAT tissue weight and gene expression. E-WAT tissue weight of LC on standard chow compared with Sham mice and SG mice that

### Impact of HFD and SG on hepatic gene expression

HFD feeding led to only slightly higher liver weight in Sham compared to LC mice, an effect that was reversed by SG (**Figure 7** a). *Hsd11b1* mRNA levels tended to increase in the Sham group, with a trend reversal after SG (**Figure 7** b). In contrast, HSD11B1 protein levels decreased under HFD, which was even more pronounced after SG. However, HSD11B1 oxoreductase activity measured in hepatic lysates showed an opposite pattern with higher activity under HFD and after SG. The intrahepatic ratios of the bile acids tauro-ursodeoxycholic acid (TUDCA) to tauro-7oxolithocholic acid (T7oxoLCA) and taurochenodeoxycholic acid (TCDCA) to T7oxoLCA were shown to be useful biomarkers of pharmacological inhibition of HSD11B1 activity (58, 59). In the present study, HFD caused a significant decrease in T7oxoLCA levels in livers of Sham and SG mice, whereby there were rather moderate effects on TUDCA and TCDCA with slightly higher levels after SG (**Figure 8**). The elevated TUDCA/T7oxoLCA and TCDCA/T7oxoLCA ratios suggest an increased HSD11B1 oxoreductase activity, in line with the higher cortisone to cortisol conversion seen in **Figure 7** b; however, this seems to be an effect of HFD and not surgical intervention, as there was no difference between Sham and SG mice (**Figure 8**).

**Figure 7:**
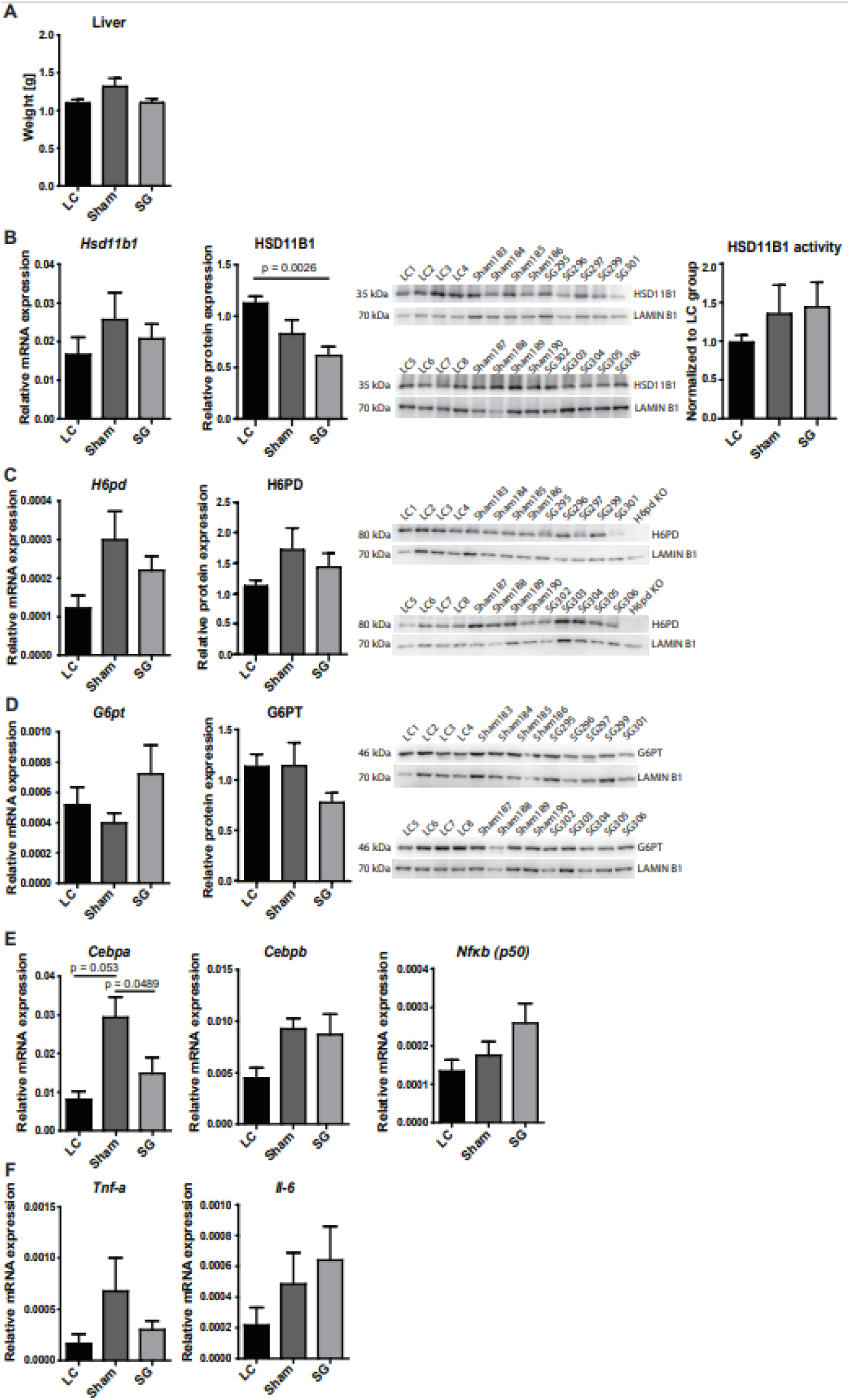
Effect of HFD and SG on hepatic gene expression. Liver tissue weight of LC (fed standard chow), Sham, and SG mice (both on HFD) (a). mRNA was quantified by RT-qPCR normalized to *18S* rRNA (a-f), and protein levels by Western blotting, normalized first to the loading control Lamin B1 and then to the expression in LC mice (b-d). HSD11B1 oxoreductase activity was measured in lysate by determining the conversion of cortisone to cortisol, followed by normalization to the activity of LC mice (b). LC1-4 served as reference samples for Sham183-186 and SG295-297, 299, 301; LC5-8 were used as reference samples for Sham187-190 and SG302-306 (b-d). Statistics: One-way ANOVA with Tukey’s multiple comparison test; p-values are indicated. Values represent mean ± SEM. LC = Lean Control (n=8, standard chow), Sham (n=8, HFD), and SG = Sleeve Gastrectomy (n=10, HFD), HFD = High-fat diet.

**Figure 8:**
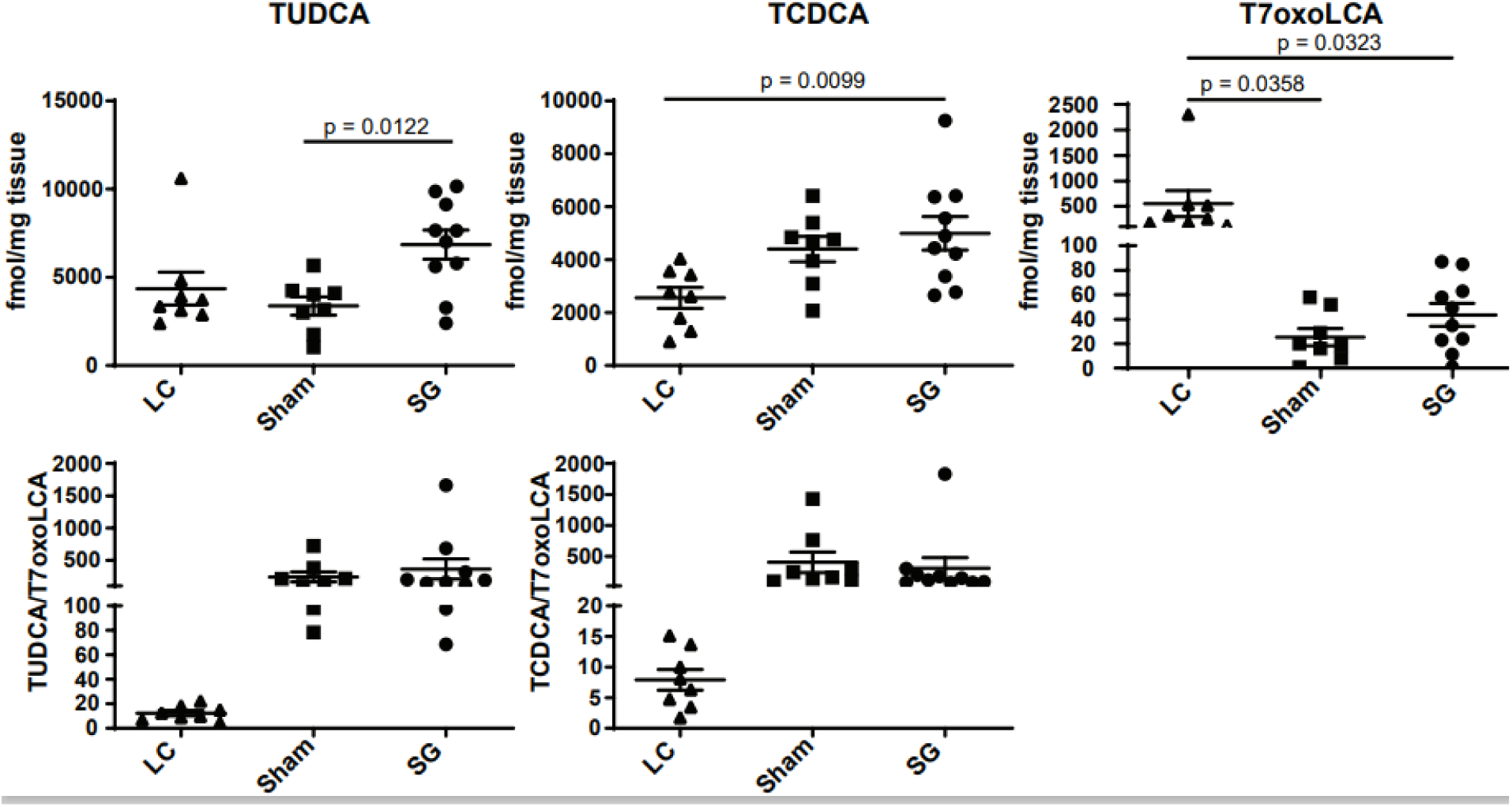
Bile acid marker for hepatic HSD11B1 activity. Intrahepatic levels of tauro-ursodeoxycholic acid (TUDCA), tauro-chenodeoxycholic acid (TCDCA), and tauro-7oxolithocholic acid (T7oxoLCA) were measured in liver tissues of LC mice on standard chow and Sham and SG mice on HFD by LC-MS/MS. The markers of HSD11B1 oxoreductase activity, *i.e.,* the ratios of TUDCA/T7oxoLCA and TCDCA/T7oxoLCA, were determined. Statistics: One-way ANOVA with Tukey’s multiple comparison test; p-values are indicated. Values represent mean ± SEM. LC = Lean Control (n=8, standard chow), Sham (n=8, HFD), and SG = Sleeve Gastrectomy (n=10, HFD), HFD = High-fat diet.

*H6pd* mRNA and H6PD protein expression both tended to be higher under HFD but were partially reversed following SG (**Figure 7** c). *G6pt* mRNA expression tended to increase, whereas its protein levels tended to decrease in the SG group (**Figure 7** d). The mRNA levels of the transcription factors *Cebpa* and *Cebpb* were higher in the Sham compared to the LC group, with a reversal for *Cebpa* but not *Cebpb* following SG, whereas *Nfκb (p50)* mRNA levels tended to be higher after SG (**Figure 7** e). *Tnf-a* mRNA tended to increase in the Sham group with a reversal after SG, whereas the pattern for *Il-6* mRNA levels resembled that of *Nfκb (p50)* (**Figure 7** f).

Next, we analyzed the expression of enzymes involved in hepatic glucocorticoid metabolism. Whereas HFD tended to increase the mRNA expression of steroid 5β-reductase (*Akr1d1*) compared to LC mice, with a more pronounced effect after SG, SG reversed this trend of higher AKR1D1 protein levels seen in Sham mice (**Figure 9** a). The mRNA and protein expression of steroid 5α-reductase (SRD5A1) tended to increase in SG mice compared to LC and Sham mice (**Figure 9** b). The mRNA levels of the glucocorticoid-metabolizing enzyme *Cyp3a11* tended to decrease under HFD, along with a significant decrease of protein levels in Sham mice compared to LC and a recovery after SG (Sham vs SG, ns, p = 0.0569) (**Figure 9** c).

**Figure 9:**
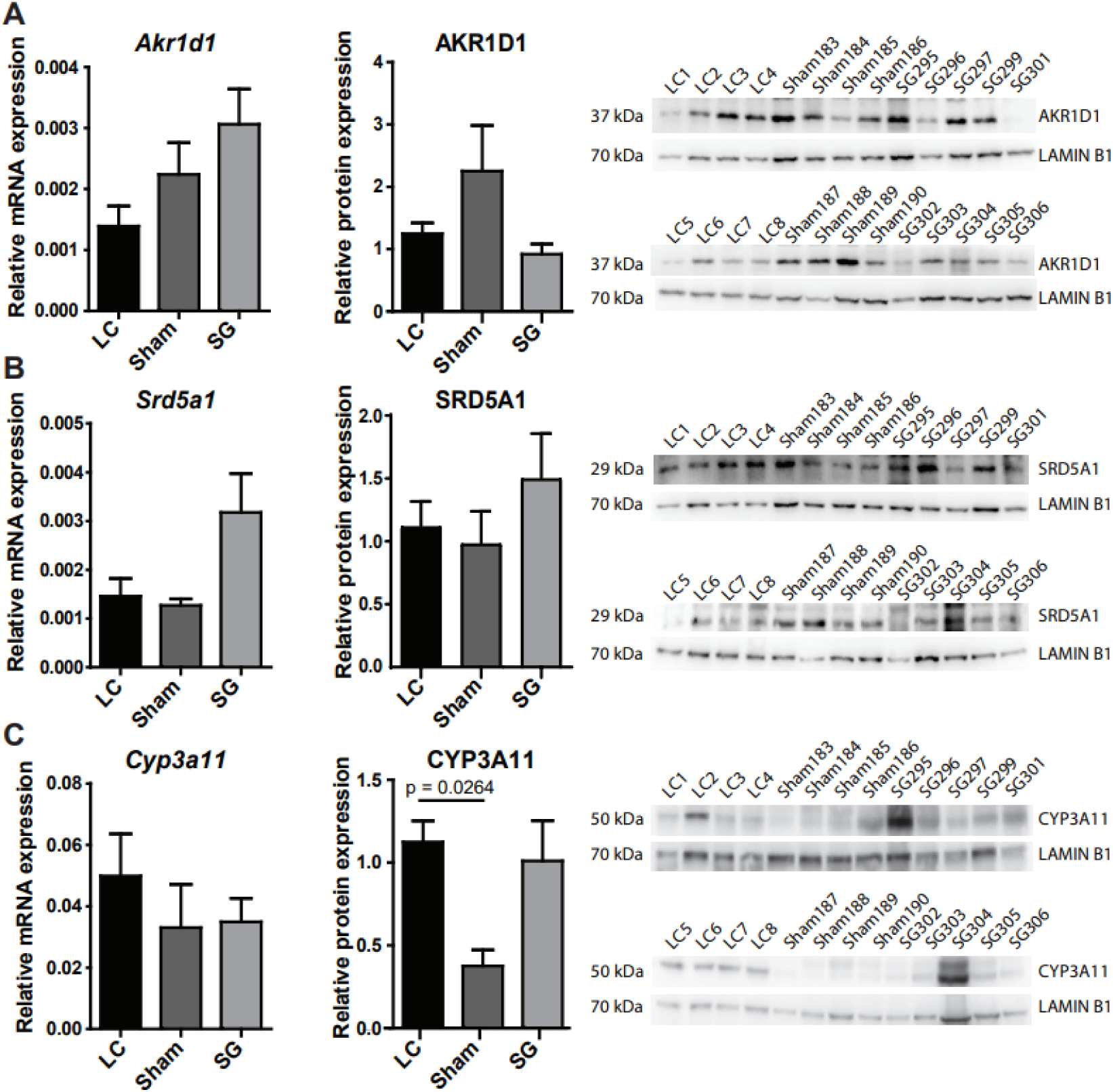
Impact of HFD and SG on enzymes involved in hepatic glucocorticoid metabolism. mRNA was measured by RT-qPCR and normalized to *18S* rRNA (a-c). Protein levels were determined by Western blotting, first normalized to the loading control Lamin B1 and then to the expression levels in LC mice. LC1-4 were used as control groups for Sham183-186 and SG295-297, 299, 301; LC5-8 served as control groups for Sham187-190 and SG302-306 (a-c). Statistical analysis was performed using one-way ANOVA with Tukey’s multiple comparison test; p-values are indicated. Values are displayed as mean ± SEM. LC = Lean Control (n=8, standard chow), Sham (n=8, HFD), and SG = Sleeve Gastrectomy (n=10, HFD), HFD = High-fat diet.

### Impact of HFD and SG on adrenal mRNA expression

The adrenal weight increased under HFD in Sham mice, with a tendency of reversal after SG (**Figure 10** a). Interestingly, the expression of the enzyme transporting cholesterol to the inner mitochondrial membrane (60), *Star*, was decreased under HFD in Sham mice, followed by at least a partial reversal after SG (**Figure 10** b). This pattern was, much less pronounced, also seen for the steroidogenic genes *Cyp11a1*, *Hsd3b1,* 11β-hydroxylase (*Cyp11b1*) and aldosterone synthase (*Cyp11b2*) (**Figure 10** b). The partial reversal of levels from Sham after SG resembled that seen for glucocorticoid serum concentrations (**Figure 4**).

**Figure 10:**
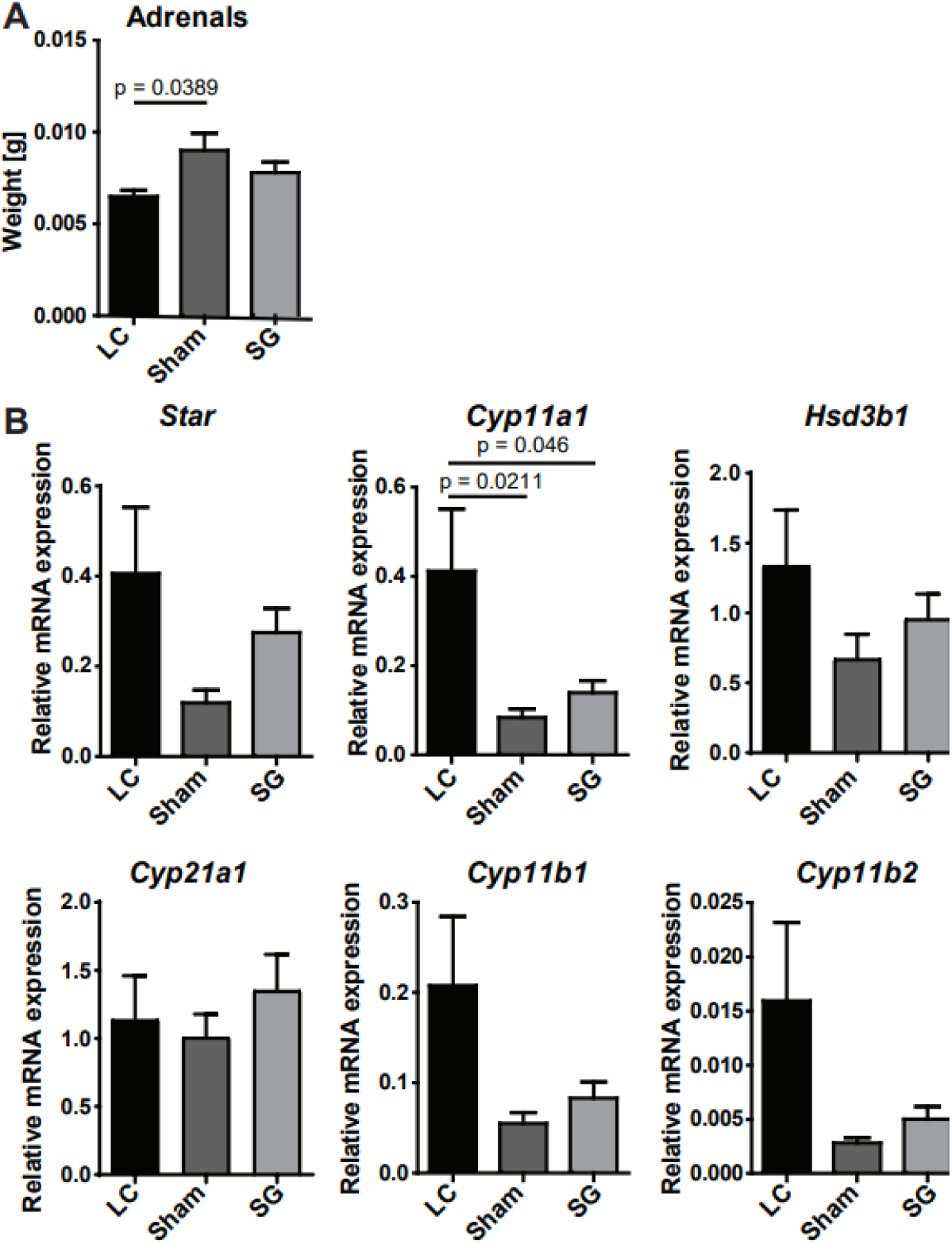
Effect of HFD and SG on adrenal tissue weight and mRNA expression. Adrenal tissue weight from LC mice on standard chow and Sham and SG mice on HFD was determined (a). mRNA was determined by RT-qPCR, normalized to *18S* rRNA (b). Statistical analysis was performed using one-way ANOVA with Tukey’s multiple comparison test; p-values are indicated. Values represent mean ± SEM. LC = Lean Control (n=8, standard chow), Sham (n=8, HFD), and SG = Sleeve Gastrectomy (n=10, HFD), HFD = High-fat diet.

## DISCUSSION

Previous studies on steroid hormonal changes following bariatric surgery have mainly focused on HSD11B1 expression in liver and adipose tissue, or investigated circulating and urinary markers of HSD11B1 and HSD11B2 activity. However, our broader understanding of how corticosteroid metabolism is altered at the systemic and tissue-specific level remains limited. This study compared sham-operated LC mice on standard chow, Sham-operated mice on HFD, and SG-treated mice on HFD (SG) to investigate changes in corticosteroid homeostasis by integrating serum and tissue-specific steroid levels, enzyme expression and activity, and expression of transcription factors and cytokines involved in the regulation of HSD11B1 and HSD11B2 expression, in order to elucidate glucocorticoid regulation in obesity and post-SG metabolic adaptations.

The reversal of the adverse effects of HFD in sham-operated mice compared to the LC group fed a standard chow on glucose metabolism and body weight seen after SG (**Figure 3**) emphasizes the relevance of our model and is in line with findings reported in human bariatric surgery studies (61). Bariatric surgery, particularly SG, is a widely used procedure and well-documented to improve metabolic parameters and improve or even resolve T2DM, a common comorbidity of obesity (62–65).

The main finding of the present study is the increase in the circulating B/A ratio in sham-operated mice on HFD compared to LC mice on standard chow and its recovery following SG. Although the recovery after SG was not statistically significant (p = 0.0567), likely due to the limited number of animals used, significant correlations of circulating B/A ratio with the OGTT at 120 min (p = 0.046) and glucose AUC (p = 0.0269) were observed when all animals were included, suggesting a beneficial impact of SG on both glucocorticoid and glucose homeostasis. This reversal in B/A ratio was likely driven by renal HSD11B2 activity, as also supported by the intrarenal B/A ratio. These findings are in line with previous experiments demonstrating a lower B/A ratio in mice undergoing OAGB surgery compared to sham-operated controls (44). Notably, the circulating F/E ratio has been reported to be lower in non-diabetic individuals compared to diabetic patients (66), reinforcing the hypothesis that the lower B/A ratio (i.e., active/inactive glucocorticoid ratio) observed in SG mice reflects a metabolic improvement.

Alterations in circulating glucocorticoid levels have been associated with diabetes and obesity (17–19, 27). In our study, serum corticosterone and 11-dehydrocorticosterone were lower in the Sham compared to the LC group, with a trend toward reversal after SG. This aligns with prior findings showing diurnal variations of corticosterone levels post-SG in mice (43). Similarly, glucocorticoid levels increased post-SG, consistent with previous observations (15, 44).

The higher *Cyp11b2* mRNA expression in LC mice likely reflects dietary differences, as the LC diet (standard chow) contained more potassium than the HFD (**Suppl. Table 5** and **Suppl. Table 6**), influencing aldosterone and 11-DOC production. Our data showed a slight decrease of mineralocorticoids post-SG, while plasma aldosterone concentrations progressively declined following RYGB surgery in patients, as well as decreased after OAGB in mice (42, 44). This more pronounced effect may vary depending on the type of surgery, the timing of measurement, and the composition of the diet. Additionally, adrenal gland hypertrophy in Sham mice, with a trend toward normalization post-SG, suggests SG may counteract obesity-associated adrenal alterations, a pattern also seen in obese Zucker rats (27).

The expression of *Hsd11b1* and *Hsd11b2* genes is regulated by transcription factors, such as Cebpa, Cebpb, and Nfκb, as well as proinflammatory cytokines, including TNF-α, IL-6 and IL-1 through their cell membrane expressed receptors (55, 67). The observed expression patterns of *Hsd11b1* and *Hsd11b2* aligned with those of their respective transcription factor dependencies, with *Hsd11b1* strongly induced by Cebpa and *Hsd11b2* by Cebpb (55), as seen for hepatic and E-WAT *Hsd11b1*, and renal *Hsd11b2* mRNA expression levels in the present study. Despite a reported reduction of cytokines in the jejunum of rats after SG (68), no significant post-SG serum cytokine changes could be observed in patients (69), suggesting either tissue-or species-specific effects. We observed an increase in renal *Tnf-a*, *Nfκb (p50)*, and *Cebpb* mRNA expression in SG mice compared to the Sham group, consistent with findings that TNF-α induces Nfκb and Cebpb in transactivation assays (70). Although TNF-α, IL-1β, and IL-6 have been shown to downregulate placental HSD11B2 (71), this effect was not observed in the kidney. Instead, an indirect regulatory mechanism involving *Nfκb (p50)* and *Cebpb* is likely. IL-1α, which increases Nfκb protein levels (72), can cooperate with *Cebpb* (70) to enhance renal HSD11B2 expression (55). Additionally, IL-1α and TNF-α can upregulate HSD11B1 through kinase-mediated Cebpb phosphorylation or via the Nfκb pathway (73), which could also influence HSD11B2 through Cebpb. Other studies regarding HSD11B2, showed elevated *HSD11B2* mRNA in adipose tissue post-surgery (48, 49). This aligns with findings that human HSD11B2 overexpression in adipocytes, prevents HFD-induced obesity in mice (74). In contrast, comparisons between severely obese and lean individuals showed an elevated urinary free E/F ratio, suggesting increased HSD11B2 activity, with an inverse association to insulin sensitivity (34), likely not only reflecting HSD11B2 but also HSD11B1 activity. This difference may be due to the influence of additional factors associated with severe obesity compared to HFD. Overall, elevated renal HSD11B2 activity in SG mice compared to Sham and LC groups (p = 0.0252) likely explains the reduction in both intrarenal and serum B/A ratios post-SG. The intrarenal B/A decrease may be less pronounced than in serum, possibly due to the presence of HSD11B1 in the renal proximal tubule, where it likely functions as an oxoreductase and influences the B/A ratio in an opposite manner than HSD11B2 (75). Isolating renal tubules and measuring enzyme activity in intact tubules could provide valuable insight into distinguishing HSD11B1 from HSD11B2 activity; however, measuring steroid levels in this context would likely not be informative, as they would probably be washed out during the isolation process. As steroids were shown to be reabsorbed mainly in the cortical collecting duct, the more pronounced effect of HSD11B2 on the circulating B/A ratio can be expected (76). These findings suggest that SG-induced HSD11B2 upregulation contributes to improved metabolic outcomes and their associated effects.

Obesity is a major risk factor for hypertension, which promotes CKD and cardiovascular complications (57, 77). The precise mechanism of obesity-induced hypertension remains unclear (57) but is associated with activation of the renin-angiotensin-aldosterone system (RAAS) and altered antihypertensive drug metabolism, leading to increased renal sodium reabsorption and impaired natriuresis (78). Patients with drug-resistant hypertension experienced higher relapse rates (27%) compared to the treatable cases (10-15%) post-SG or OAGB (79), with similar relapse rates observed for SG and RYGB (10-14%) (80). Mineralocorticoid receptor (MR) activation in sodium-transporting epithelial cells is essential for blood pressure regulation, driving sodium and water retention while promoting potassium excretion. Glucocorticoids can also bind to the MR with an affinity comparable to that of mineralocorticoids (81–83). HSD11B2 is co-localized with the MR in mineralocorticoid target tissues and prevents glucocorticoid-mediated MR activation by converting active glucocorticoids into their inactive forms, thereby maintaining sodium homeostasis and normotension (84). The increase of renal HSD11B2 post-SG is likely a protective mechanism against obesity-induced hypertension to limit glucocorticoid-mediated MR activation. This proposes beneficial effects of SG *via* renal HSD11B2 in obesity, associated CKD, and metabolic diseases. MR antagonists effectively lower resistant hypertension even in the absence of altered aldosterone levels, whereas aldosterone synthesis inhibitors show limited efficacy in this context (85–87), further emphasizing the importance of glucocorticoid-mediated MR activation. In addition, obesity is associated with reduced renal HSD11B2, as demonstrated in obese Zucker rats and dogs (27, 77), while *Hsd11b2* knockout mice develop salt-sensitive hypertension (88). Moreover, CKD patients exhibit an elevated circulating F/E ratio, indicative of reduced HSD11B2 activity (89). Interestingly, renal HSD11B2 activity was not different between LC and Sham mice, which may be due to higher potassium intake in LC mice. Potassium increases aldosterone synthesis via CYP11B2 (90), a pattern reflected in our study, suggesting a diminished activation of the MR by corticosterone in LC mice. Future studies should assess 24 h urinary F/E ratios in patients before and after SG to determine the utility of the F/E ratio as an informative marker for reduced glucocorticoid-dependent MR activation and hypertension. Moreover, therapeutic strategies to enhance renal HSD11B2 activity should be explored.

Our results suggest that hepatic HSD11B1 activity did not substantially contribute to the circulating B/A ratio. The intrahepatic B/A ratio remained unchanged across groups and showed no correlation with OGTT outcomes. This was supported by HSD11B1 activity data and the bile acid marker of HSD11B1 activity measured in liver tissue (58, 59). *Hsd11b1* and *Cebpa* mRNA expression were closely linked in our data, consistent with reports of Cebpa-driven HSD11B1 activation in liver (55, 91). Although it was reported that TNF-α and IL-1 regulate HSD11B1 via Cebpb and Nfκb (67, 73, 92), our data suggest a more complex interaction, where *Il-6* and *Nfκb (p50)* mRNA, rather than *Tnf-a* and *Cebpa* mRNA expression, correlated with HSD11B1 activity. Unlike previous reports of decreased hepatic HSD11B1 activity in obese Zucker rats (27), our study found no significant changes in hepatic HSD11B1 protein or activity, aligning with findings that hepatic glucocorticoid regeneration remains localized (93). However, strain-specific differences may also need to be considered. Bile acid measurements (TUDCA, TCDCA, T7oxoLCA) suggested microbiota-driven changes in HFD-fed mice, with decreased T7oxoLCA under HFD. A recent study reported that obese patients showed reduced microbial diversity and lower levels of secondary bile acids, both of which were gradually recovered over time following SG (94), potentially explaining the lack of reversal observed in our findings due to the time-dependent nature of these changes. Even though H6PD, required for HSD11B1 oxoreductase function, depends on glucose metabolism through G6PT (7, 95), findings in db/db (obese and diabetic) mice showing increased gene expression of hepatic *H6pd* and *G6pt* compared to a lean control (96) were moderately reflected for H6PD and G6PT in our data. In conclusion, no substantial changes in HSD11B1 expression or activity could be observed, supported by the stable intrahepatic B/A ratio. Investigating hepatic corticosteroid metabolism, *5α-reductase* expression has been linked to fasting serum insulin levels in obese patients (33), and post-RYGB reductions in urinary 5α-THF/5β-THF ratios suggest decreased 5α-reductase-mediated corticosteroid metabolism (47), findings differing from our observations. Our data on corticosteroid metabolism seemed to shift post-SG to increased 5α-reductase (SRD5A1), likely compensating for decreased 5β-reductase (AKR1D1). Elevated AKR1D1 in the Sham group aligns with findings that urinary 5β-THE is elevated in obesity (97) and 5β-THF/F and 5β-THE/E ratios correlate with liver fat (30). Additionally, a marker of CYP3A4 (human homolog of CYP3A11) was decreased in hypertensive children (98), aligning with lower CYP3A11 in Sham mice in our study.

With respect to adipose tissue, we found that SG normalized E-WAT weight, reduced *Fasn* expression associated with HFD-induced adipogenesis, and restored elevated *Lipe* (HSL) expression, consistent with earlier findings and reduced HSL activity post-RYGB (99–101). HSD11B1 upregulation in adipose tissue in obesity and its contribution to weight gain in mice have been well-documented (25, 28). We found *Cebpa* mRNA expression to be linked with *Hsd11b1* mRNA expression, as previously reported (55, 102). However, on the protein level, our findings regarding HSD11B1 contrasted with those in the literature. Whilst VAT HSD11B1 activity was previously reported to increase in obesity (27), we found lower HSD11B1 protein expression in E-WAT of Sham mice, and recovery following SG. Interestingly, Morton *et al.* reported that chronic HFD feeding led to HSD11B1 downregulation in adipose tissue (e.g., subcutaneous, epididymal) across various mouse strains, suggesting a compensatory reduction in glucocorticoid signaling after HFD (32), consistent with our observations. Thus, an explanation may be that HFD induces an adaptive, protective response, while in chronic obesity, particularly in a low-grade inflammatory environment, *Hsd11b1* expression may be upregulated by proinflammatory cytokines and Nfκb in an attempt to lower local inflammation. Direct HSD11B1 enzyme activity measurements were not possible due to insufficient amounts of fat tissue from LC and SG mice. *H6PD* expression has been associated with metabolic syndrome and *HSD11B1* mRNA levels (103), and both *H6pd* and *G6pt* expression levels were reported to be elevated in adipose tissue of db/db mice compared to lean controls (96). H6PD and G6PT expression levels and their relationship with obesity were only very moderately associated in our study. Lower H6PD and G6PT expression in LC and SG mice may confer metabolic benefits, as adipose tissue-specific H6PD knockout mice showed improved insulin sensitivity and reduced visceral fat (104).

Limitations of our study include the use of male mice only, dietary differences between the LC and HFD-fed groups, a relatively low number of mice per group, and, for steroid analysis, the lack of 24-hour urine collections, issues clearly requiring consideration in future studies.

In conclusion, our study demonstrated that SG normalized glucocorticoid homeostasis in obese mice under HFD, highlighting the complex regulation of these hormones in metabolic disease, influenced by both dietary composition and disease state (HFD vs. low-grade inflammation in obesity). The correlation between the circulating B/A ratio and OGTT and glucose AUC outcomes proposes the B/A ratio as a useful marker for metabolic improvement following SG in rodents and the F/E ratio as the corresponding marker in humans. The increased renal HSD11B2 activity and reduced B/A ratios in kidney and serum following SG suggest beneficial effects in metabolic diseases, CKD, and hypertension. Further research is needed to better understand the contribution of HSD11B1 function, also considering the role of H6PD in its oxoreductase activity. Furthermore, the F/E ratio should be investigated in patients after bariatric surgery as an informative marker for glucocorticoid-mediated MR activation, and strategies to enhance renal HSD11B2 activity for clinical application should be explored.

## DATA AVAILABILITY

The datasets supporting the conclusions of this article are included in the article and its additional files. The data generated during and/or analyzed during the current study are available at Zenodo (link will be included in the final version).

## SUPPLEMENTAL MATERIAL

Supplemental Figures 1-2, Supplemental Tables 1-7.

## GLOSSARY

A: 11-Dehydrocorticosterone
ACTH: Adrenocorticotropic hormone
AUC: Area under the curve
B: Corticosterone
Cebp: CCAAT enhancer binding protein
CKD: Chronic kidney disease
CRH: Corticotrophin-releasing hormone
E: Cortisone
ER: Endoplasmic reticulum
E-WAT: Epididymal white adipose tissue
F: Cortisol
H6PD: Hexose-6-phosphate dehydrogenase
HFD: High-fat diet
HPA: Hypothalamic-pituitary-adrenal
HSD11B: 11β-hydroxysteroid dehydrogenase
IL: Interleukin
ISTD: Internal standards
LC-MS/MS: Liquid chromatography tandem mass spectrometry
LLE: Liquid-liquid-extraction
MR: Mineralocorticoid receptor
NFκB: Nuclear factor kappa B
OAGB: One-anastomosis gastric bypass
OGTT: Oral glucose tolerance test
PBS: Phosphate-buffered saline
RAAS: Renin-angiotensin-aldosterone system
RYGB: Roux-en-Y gastric bypass
SAT: Subcutaneous adipose tissue
SEM: Standard error of the mean
SG: Sleeve gastrectomy
SPE: Solid-phase extraction
TCDCA: Tauro-chenodeoxycholic acid
T2DM: Type 2 diabetes mellitus
THE: Tetrahydrocortisone
THF: Tetrahydrocortisol
TNF-α: Tumor-necrosis factor alpha
T7oxoLCA: Tauro-7oxolithocholic acid
TUDCA: Tauro-ursodeoxycholic acid
VAT: Visceral adipose tissue

## ACKNOWLEDGMENTS

We thank Prof. Carsten Gründemann for support with the CytoFLEX S Flow Cytometer to measure renal cytokines. ChatGPT, powered by OpenAI, running with GPT-4.5 architecture, was used to paraphrase and rewrite paragraphs. The tool was used in a manner that does not conflict with APS ethical policies and the authors take full responsibility for the content.

## DISCLAIMERS

The authors have nothing to disclaim.

## AUTHOR CONTRIBUTIONS

SOM, AM, AO, and EGS conceived and designed research;

SOM, AM, CFR, DVW, FLJ, and CG performed experiments and analyzed data;

SOM and AO interpreted the results of experiments;

SOM prepared figures and drafted the manuscript;

SOM, AM, CFR, AO, EGS, DVW, FLJ, and CG edited, revised the manuscript, and approved the final version of the manuscript.

## FIGURE LEGENDS

**Suppl. Figure 1:**
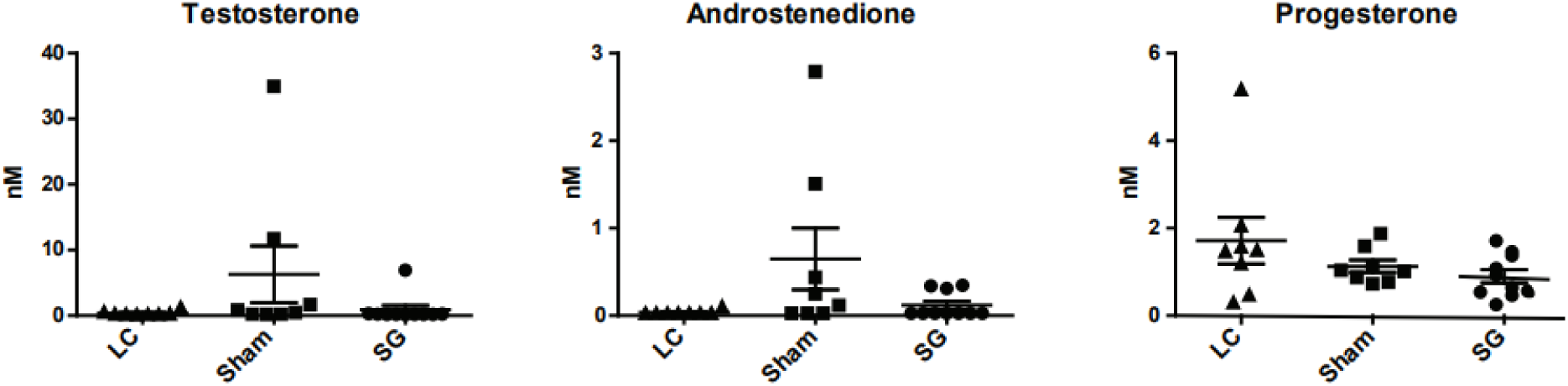
Impact of HFD and SG on sex steroid hormone levels. Steroid hormone levels were determined by LC-MS/MS. Statistical analysis was performed using one-way ANOVA with Tukey’s multiple comparison test; p-values are indicated. Values represent mean ± SEM. LC = Lean Control (n=8, standard chow), Sham (n=8, HFD), and SG = Sleeve Gastrectomy (n=10, HFD), HFD = High-fat diet.

**Suppl. Figure 2:**
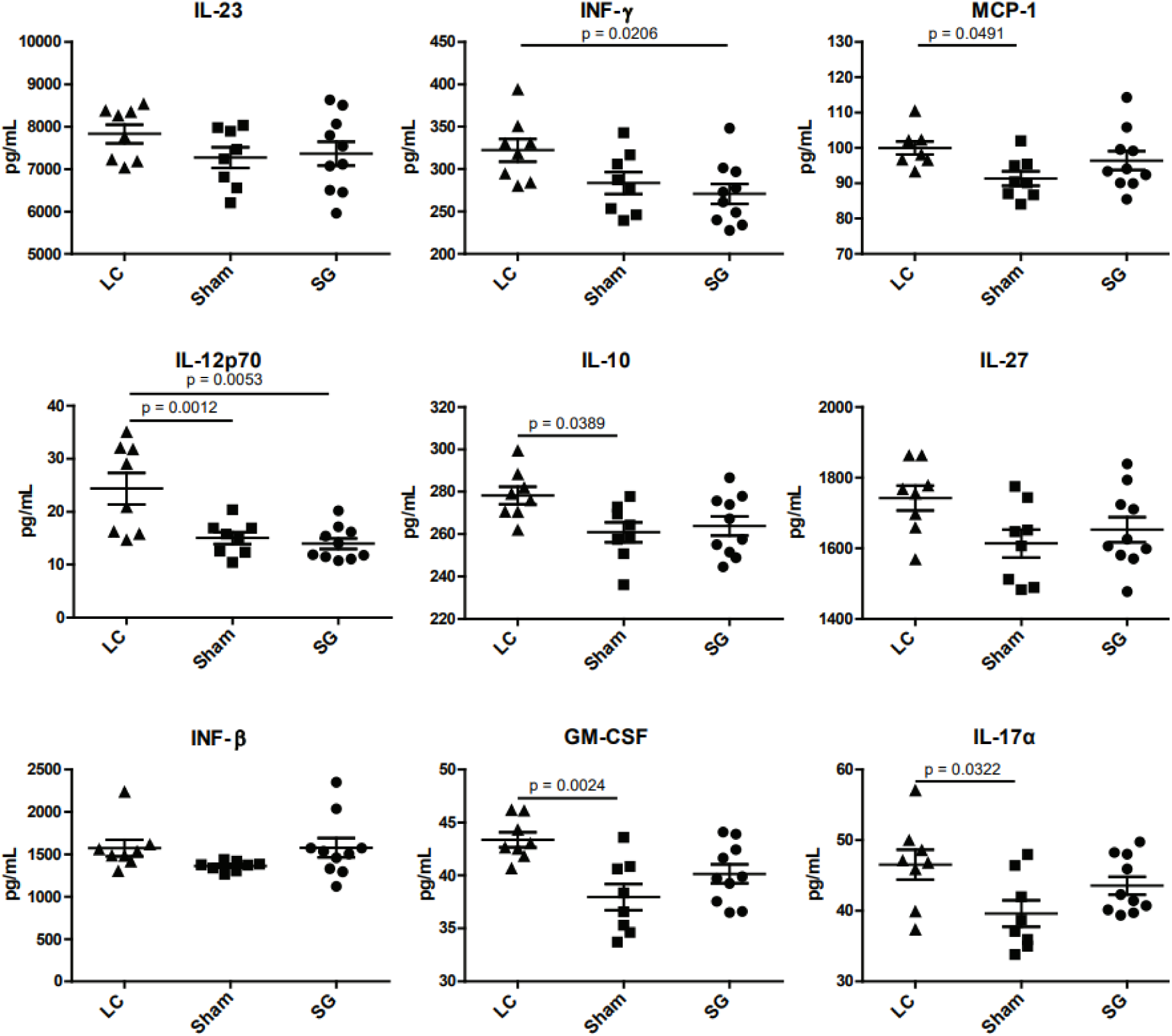
Effect of HFD and SG on intrarenal cytokine levels. Cytokine **p**rotein levels were detected by flow cytometry (LEGENDplex™ Mouse Inflammation Panel (13-plex)). Statistical analysis was performed using one-way ANOVA with Tukey’s multiple comparison test; p-values are indicated. Values represent mean ± SEM. LC = Lean Control (n=8, standard chow), Sham (n=8, HFD), and SG = Sleeve Gastrectomy (n=10, HFD), HFD = High-fat diet.

## Supplemental Tables

**Suppl. Table 1:**
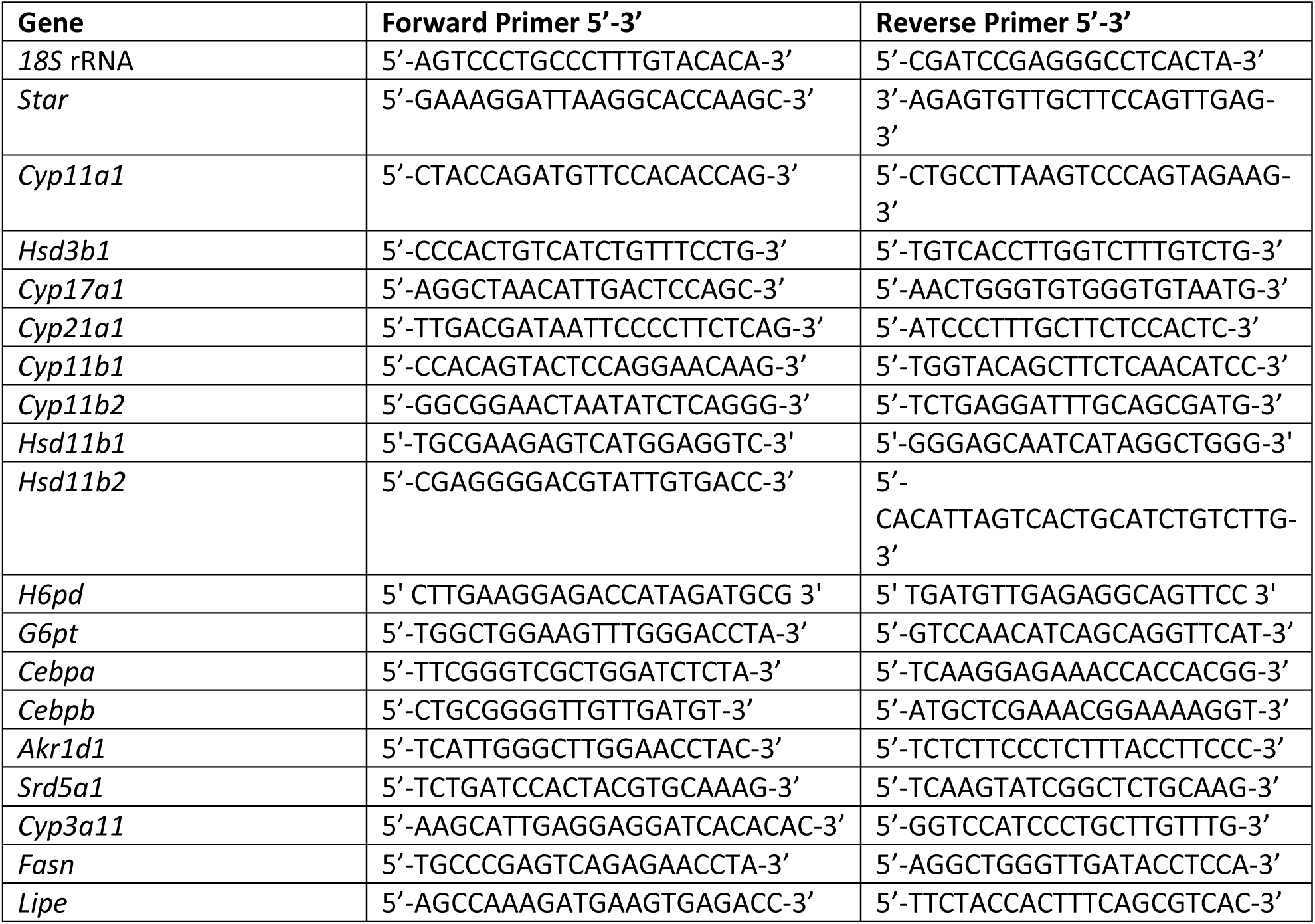
Gene and corresponding primer sequences for RT-qPCR.

**Suppl. Table 2:** Antibodies used to assess liver homogenates for Western blotting and respective dilutions.

| Protein | Company | Distributor | REF nr. | Primary antibody dilution | Secondary antibody dilution |
| --- | --- | --- | --- | --- | --- |
| Lamin B1 | Abcam | Lucerna Chem | ab133741 | 1:1000 | 1:5000 |
| HSD11B1 | Cayman Chemicals | Chemie Brunschwig | 10004303 | 1:5000 | 1:5000 |
| H6PD | Sigma-Aldrich | Sigma-Aldrich | HPA004824 | 1:1000 | 1:2000 |
| G6PT | Proteintech | LubioScience | 20612-1-AP | 1:2000 | 1:5000 |
| SRD5A1 | Proteintech | LubioScience | 66329-1-Ig | 1:1000 | 1:2000 |
| AKR1D1 | SantaCruz | LabForce | sc-365932 | 1:500 | 1:2000 |
| CYP3A1 | SantaCruz | LabForce | sc-53246 | 1:1000 | 1:2000 |

**Suppl. Table 3:** Antibodies used to assess epididymal white adipose tissue (E-WAT) homogenates for Western blotting and respective dilutions.

| Protein | Company | Distributor | REF nr. | Primary antibody dilution | Secondary antibody dilution |
| --- | --- | --- | --- | --- | --- |
| β-Actin (ACTB) | SantaCruz | LabForce | sc-47778 | 1:2000 | 1:2000 |
| HSD11B1 | Cayman Chemicals | Chemie Brunschwig | 10004303 | 1:2000 | 1:2000 |
| H6PD | Sigma-Aldrich | Sigma-Aldrich | HPA004824 | 1:500 | 1:1000 |
| G6PT | Proteintech | LubioScience | 20612-1-AP | 1:2000 | 1:2000 |

**Suppl. Table 4:** Secondary antibodies used for Western blotting.

| Secondary antibodies | Company | Distributor | REF nr. |
| --- | --- | --- | --- |
| Rabbit | Cell Signaling | BioConcept | 7074S |
| Mouse | Sigma-Aldrich | Sigma-Aldrich | A0168 |

**Suppl. Table 5:** Standard chow food composition (Pico5053; Laboratory Diet, St. Louis, MO).

| Nutrition's | % in food |
| --- | --- |
| Protein | 20 |
| Carbohydrates | 57.6 |
| Fat (total) | 11.6 |
| Minerals (total) | 6.1 |
| Calcium | 0.81 |
| Phosphorus | 0.96 |
| Potassium | 1.07 |
| Sodium | 0.3 |
| Magnesium | 0.22 |
| Sulfur | 0.34 |
| Chlorine | 0.51 |
| Vitamin Mix | Small amounts |

**Suppl. Table 6:**
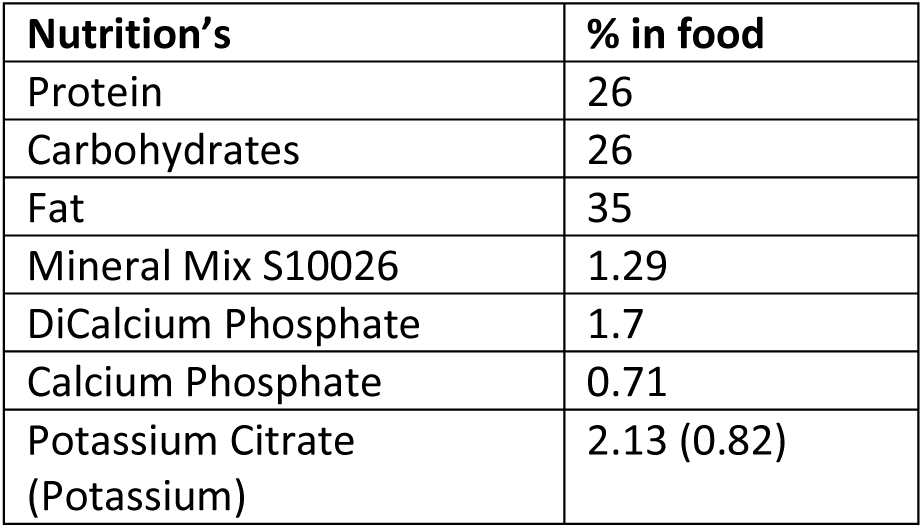
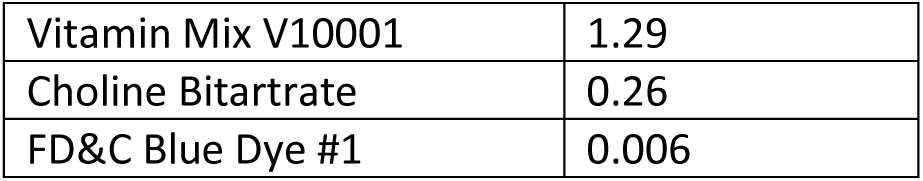
High-fat diet composition (D12492; Research Diets, New Brunswick, NJ).

| Nutrition's | % in food |
| --- | --- |
| Protein | 26 |
| Carbohydrates | 26 |
| Fat | 35 |
| Mineral Mix S10026 | 1.29 |
| DiCalcium Phosphate | 1.7 |
| Calcium Phosphate | 0.71 |
| Potassium Citrate (Potassium) | 2.13 (0.82) |
| Vitamin Mix V10001 | 1.29 |
| Choline Bitartrate | 0.26 |
| FD&C Blue Dye #1 | 0.006 |

**Suppl. Table 7:** Post-surgery gel diet composition (DietGel® Recovery, ClearH2O, Westbrook, ME).

| **Nutrition's** | **% in food** |
| --- | --- |
| Protein | 0.6 |
| Carbohydrates | 23.2 |
| Fat | 1.9 |
| Moisture | 71-75 |
| Calcium | 0.035 |
| Phosphorus | 0.048 |
| Potassium | 0.086 |
| Sodium | 0.011 |
| Magnesium | 0.0003 |
| Chloride | 0.115 |

## REFERENCES

1. Gathercole LL, Lavery GG, Morgan SA, Cooper MS, Sinclair AJ, Tomlinson JW, et al. 11beta-Hydroxysteroid dehydrogenase 1: translational and therapeutic aspects. Endocr Rev. 2013;34(4):525–55.

2. Fadel L, Dacic M, Fonda V, Sokolsky BA, Quagliarini F, Rogatsky I, et al. Modulating glucocorticoid receptor actions in physiology and pathology: Insights from coregulators. Pharmacol Ther. 2023;251:108531.

3. Gjerstad JK, Lightman SL, Spiga F. Role of glucocorticoid negative feedback in the regulation of HPA axis pulsatility. Stress. 2018;21(5):403–16.

4. Miller WL, Auchus RJ. The molecular biology, biochemistry, and physiology of human steroidogenesis and its disorders. Endocr Rev. 2011;32(1):81–151.

5. Odermatt A, Klusonova P. 11beta-Hydroxysteroid dehydrogenase 1: Regeneration of active glucocorticoids is only part of the story. J Steroid Biochem Mol Biol. 2015;151:85–92.

6. Odermatt A, Kratschmar DV. Tissue-specific modulation of mineralocorticoid receptor function by 11beta-hydroxysteroid dehydrogenases: an overview. Mol Cell Endocrinol. 2012;350(2):168–86.

7. Atanasov AG, Nashev LG, Schweizer RA, Frick C, Odermatt A. Hexose-6-phosphate dehydrogenase determines the reaction direction of 11beta-hydroxysteroid dehydrogenase type 1 as an oxoreductase. FEBS Lett. 2004;571(1-3):129–33.

8. Banhegyi G, Benedetti A, Fulceri R, Senesi S. Cooperativity between 11beta-hydroxysteroid dehydrogenase type 1 and hexose-6-phosphate dehydrogenase in the lumen of the endoplasmic reticulum. J Biol Chem. 2004;279(26):27017–21.

9. Odermatt A, Arnold P, Stauffer A, Frey BM, Frey FJ. The N-terminal anchor sequences of 11beta-hydroxysteroid dehydrogenases determine their orientation in the endoplasmic reticulum membrane. J Biol Chem. 1999;274(40):28762–70.

10. Livingstone DE, Barat P, Di Rollo EM, Rees GA, Weldin BA, Rog-Zielinska EA, et al. 5alpha-Reductase type 1 deficiency or inhibition predisposes to insulin resistance, hepatic steatosis, and liver fibrosis in rodents. Diabetes. 2015;64(2):447–58.

11. Palermo M, Marazzi MG, Hughes BA, Stewart PM, Clayton PT, Shackleton CH. Human Delta4-3-oxosteroid 5beta-reductase (AKR1D1) deficiency and steroid metabolism. Steroids. 2008;73(4):417–23.

12. Galteau MM, Shamsa F. Urinary 6beta-hydroxycortisol: a validated test for evaluating drug induction or drug inhibition mediated through CYP3A in humans and in animals. Eur J Clin Pharmacol. 2003;59(10):713–33.

13. Kivimaki M, Strandberg T, Pentti J, Nyberg ST, Frank P, Jokela M, et al. Body-mass index and risk of obesity-related complex multimorbidity: an observational multicohort study. Lancet Diabetes Endocrinol. 2022;10(4):253–63.

14. Thomas MC. Type 2 Diabetes and Heart Failure: Challenges and Solutions. Curr Cardiol Rev. 2016;12(3):249–55.

15. Woods CP, Corrigan M, Gathercole L, Taylor A, Hughes B, Gaoatswe G, et al. Tissue specific regulation of glucocorticoids in severe obesity and the response to significant weight loss following bariatric surgery (BARICORT). J Clin Endocrinol Metab. 2015;100(4):1434–44.

16. Anagnostis P, Athyros VG, Tziomalos K, Karagiannis A, Mikhailidis DP. Clinical review: The pathogenetic role of cortisol in the metabolic syndrome: a hypothesis. J Clin Endocrinol Metab. 2009;94(8):2692–701.

17. Rosmond R, Dallman MF, Bjorntorp P. Stress-related cortisol secretion in men: relationships with abdominal obesity and endocrine, metabolic and hemodynamic abnormalities. J Clin Endocrinol Metab. 1998;83(6):1853–9.

18. Marin P, Darin N, Amemiya T, Andersson B, Jern S, Bjorntorp P. Cortisol secretion in relation to body fat distribution in obese premenopausal women. Metabolism. 1992;41(8):882–6.

19. Epel ES, McEwen B, Seeman T, Matthews K, Castellazzo G, Brownell KD, et al. Stress and body shape: stress-induced cortisol secretion is consistently greater among women with central fat. Psychosom Med. 2000;62(5):623–32.

20. Andrew R, Gale CR, Walker BR, Seckl JR, Martyn CN. Glucocorticoid metabolism and the Metabolic Syndrome: associations in an elderly cohort. Exp Clin Endocrinol Diabetes. 2002;110(6):284–90.

21. Petersen KF, Shulman GI. Etiology of insulin resistance. Am J Med. 2006;119(5 Suppl 1):S10–6.

22. Samuel VT, Petersen KF, Shulman GI. Lipid-induced insulin resistance: unravelling the mechanism. Lancet. 2010;375(9733):2267–77.

23. Dourakis SP, Sevastianos VA, Kaliopi P. Acute severe steatohepatitis related to prednisolone therapy. Am J Gastroenterol. 2002;97(4):1074–5.

24. Ehrhart-Bornstein M, Lamounier-Zepter V, Schraven A, Langenbach J, Willenberg HS, Barthel A, et al. Human adipocytes secrete mineralocorticoid-releasing factors. Proc Natl Acad Sci U S A. 2003;100(24):14211–6.

25. Masuzaki H, Paterson J, Shinyama H, Morton NM, Mullins JJ, Seckl JR, et al. A transgenic model of visceral obesity and the metabolic syndrome. Science. 2001;294(5549):2166–70.

26. Paterson JM, Morton NM, Fievet C, Kenyon CJ, Holmes MC, Staels B, et al. Metabolic syndrome without obesity: Hepatic overexpression of 11beta-hydroxysteroid dehydrogenase type 1 in transgenic mice. Proc Natl Acad Sci U S A. 2004;101(18):7088–93.

27. Livingstone DE, Jones GC, Smith K, Jamieson PM, Andrew R, Kenyon CJ, et al. Understanding the role of glucocorticoids in obesity: tissue-specific alterations of corticosterone metabolism in obese Zucker rats. Endocrinology. 2000;141(2):560–3.

28. Desbriere R, Vuaroqueaux V, Achard V, Boullu-Ciocca S, Labuhn M, Dutour A, et al. 11beta-hydroxysteroid dehydrogenase type 1 mRNA is increased in both visceral and subcutaneous adipose tissue of obese patients. Obesity (Silver Spring). 2006;14(5):794–8.

29. Munoz R, Carvajal C, Escalona A, Boza C, Perez G, Ibanez L, et al. 11beta-hydroxysteroid dehydrogenase type 1 is overexpressed in subcutaneous adipose tissue of morbidly obese patients. Obes Surg. 2009;19(6):764–70.

30. Westerbacka J, Yki-Jarvinen H, Vehkavaara S, Hakkinen AM, Andrew R, Wake DJ, et al. Body fat distribution and cortisol metabolism in healthy men: enhanced 5beta-reductase and lower cortisol/cortisone metabolite ratios in men with fatty liver. J Clin Endocrinol Metab. 2003;88(10):4924–31.

31. Rask E, Olsson T, Soderberg S, Andrew R, Livingstone DE, Johnson O, et al. Tissue-specific dysregulation of cortisol metabolism in human obesity. J Clin Endocrinol Metab. 2001;86(3):1418–21.

32. Morton NM, Ramage L, Seckl JR. Down-regulation of adipose 11beta-hydroxysteroid dehydrogenase type 1 by high-fat feeding in mice: a potential adaptive mechanism counteracting metabolic disease. Endocrinology. 2004;145(6):2707–12.

33. Baudrand R, Dominguez JM, Carvajal CA, Riquelme A, Campino C, Macchiavello S, et al. Overexpression of hepatic 5alpha-reductase and 11beta-hydroxysteroid dehydrogenase type 1 in visceral adipose tissue is associated with hyperinsulinemia in morbidly obese patients. Metabolism. 2011;60(12):1775–80.

34. Mussig K, Remer T, Haupt A, Gallwitz B, Fritsche A, Haring HU, et al. 11beta-hydroxysteroid dehydrogenase 2 activity is elevated in severe obesity and negatively associated with insulin sensitivity. Obesity (Silver Spring). 2008;16(6):1256–60.

35. Salas-Parra RD, Smolkin C, Choksi S, Pryor AD. Bariatric Surgery: Current Trends and Newer Surgeries. Gastrointest Endosc Clin N Am. 2024;34(4):609–26.

36. Leyaro B, Boakye D, Howie L, Ali A, Carragher R. Associations between Type of Bariatric Surgery and Obstructive Sleep Apnoea, Employment Outcomes, and Body Image Satisfaction: A Systematic Review and Meta-Analysis. Obes Facts. 2024:1–15.

37. Wilson RB, Lathigara D, Kaushal D. Systematic Review and Meta-Analysis of the Impact of Bariatric Surgery on Future Cancer Risk. Int J Mol Sci. 2023;24(7).

38. Robert M, Espalieu P, Pelascini E, Caiazzo R, Sterkers A, Khamphommala L, et al. Efficacy and safety of one anastomosis gastric bypass versus Roux-en-Y gastric bypass for obesity (YOMEGA): a multicentre, randomised, open-label, non-inferiority trial. Lancet. 2019;393(10178):1299–309.

39. Lee WJ, Yu PJ, Wang W, Chen TC, Wei PL, Huang MT. Laparoscopic Roux-en-Y versus mini-gastric bypass for the treatment of morbid obesity: a prospective randomized controlled clinical trial. Ann Surg. 2005;242(1):20–8.

40. Rubino F, Gagner M, Gentileschi P, Kini S, Fukuyama S, Feng J, et al. The early effect of the Roux-en-Y gastric bypass on hormones involved in body weight regulation and glucose metabolism. Ann Surg. 2004;240(2):236–42.

41. Ruiz-Tovar J, Oller I, Galindo I, Llavero C, Arroyo A, Calero A, et al. Change in levels of C-reactive protein (CRP) and serum cortisol in morbidly obese patients after laparoscopic sleeve gastrectomy. Obes Surg. 2013;23(6):764–9.

42. Berney M, Vakilzadeh N, Maillard M, Faouzi M, Grouzmann E, Bonny O, et al. Bariatric Surgery Induces a Differential Effect on Plasma Aldosterone in Comparison to Dietary Advice Alone. Front Endocrinol (Lausanne). 2021;12:745045.

43. Akalestou E, Lopez-Noriega L, Christakis I, Hu M, Miras AD, Leclerc I, et al. Vertical sleeve gastrectomy normalizes circulating glucocorticoid levels and lowers glucocorticoid action tissue-selectively in mice. Front Endocrinol (Lausanne). 2022;13:1020576.

44. Lazaridis, II, Bosch AJT, Keller L, Low AJY, Tamarelle J, Moser SO, et al. Metabolic outcomes in obese mice undergoing one-anastomosis gastric bypass (OAGB) with a long or a short biliopancreatic limb. Am J Physiol Endocrinol Metab. 2024;326(6):E819–E31.

45. Leyvraz C, Verdumo C, Suter M, Paroz A, Calmes JM, Marques-Vidal PM, et al. Changes in gene expression profile in human subcutaneous adipose tissue during significant weight loss. Obes Facts. 2012;5(3):440–51.

46. Simonyte K, Olsson T, Naslund I, Angelhed JE, Lonn L, Mattsson C, et al. Weight loss after gastric bypass surgery in women is followed by a metabolically favorable decrease in 11beta-hydroxysteroid dehydrogenase 1 expression in subcutaneous adipose tissue. J Clin Endocrinol Metab. 2010;95(7):3527–31.

47. Rask E, Simonyte K, Lonn L, Axelson M. Cortisol metabolism after weight loss: associations with 11 beta-HSD type 1 and markers of obesity in women. Clin Endocrinol (Oxf). 2013;78(5):700–5.

48. Methlie P, Dankel S, Myhra T, Christensen B, Gjerde J, Fadnes D, et al. Changes in adipose glucocorticoid metabolism before and after bariatric surgery assessed by direct hormone measurements. Obesity (Silver Spring). 2013;21(12):2495–503.

49. Podraza J, Gutowska K, Lenartowicz A, Wasowski M, Jonas MI, Bartoszewicz Z, et al. The Role of microRNA in the Regulation of Cortisol Metabolism in the Adipose Tissue in the Course of Obesity. Int J Mol Sci. 2024;25(10).

50. Kley M, Moser SO, Winter DV, Odermatt A. In vitro methods to assess 11beta-hydroxysteroid dehydrogenase type 1 activity. Methods Enzymol. 2023;689:121–65.

51. Kley M, Moser SO, Winter DV, Odermatt A. In vitro methods to assess 11beta-hydroxysteroid dehydrogenase type 2 activity. Methods Enzymol. 2023;689:167–200.

52. Strajhar P, Schmid Y, Liakoni E, Dolder PC, Rentsch KM, Kratschmar DV, et al. Acute Effects of Lysergic Acid Diethylamide on Circulating Steroid Levels in Healthy Subjects. J Neuroendocrinol. 2016;28(3):12374.

53. Gomez C, Stucheli S, Kratschmar DV, Bouitbir J, Odermatt A. Development and Validation of a Highly Sensitive LC-MS/MS Method for the Analysis of Bile Acids in Serum, Plasma, and Liver Tissue Samples. Metabolites. 2020;10(7).

54. Hamid AK, Pastor Arroyo EM, Calvet C, Hewitson TD, Muscalu ML, Schnitzbauer U, et al. Phosphate Restriction Prevents Metabolic Acidosis and Curbs Rise in FGF23 and Mortality in Murine Folic Acid-Induced AKI. J Am Soc Nephrol. 2024;35(3):261–80.

55. Balazs Z, Schweizer RA, Frey FJ, Rohner-Jeanrenaud F, Odermatt A. DHEA induces 11 –HSD2 by acting on CCAAT/enhancer-binding proteins. J Am Soc Nephrol. 2008;19(1):92–101.

56. Kostadinova RM, Nawrocki AR, Frey FJ, Frey BM. Tumor necrosis factor alpha and phorbol 12-myristate-13-acetate down-regulate human 11beta-hydroxysteroid dehydrogenase type 2 through p50/p50 NF-kappaB homodimers and Egr-1. FASEB J. 2005;19(6):650–2.

57. Hall JE, do Carmo JM, da Silva AA, Wang Z, Hall ME. Obesity, kidney dysfunction and hypertension: mechanistic links. Nat Rev Nephrol. 2019;15(6):367–85.

58. Weingartner M, Stucheli S, Kratschmar DV, Birk J, Klusonova P, Chapman KE, et al. The ratio of ursodeoxycholyltaurine to 7-oxolithocholyltaurine serves as a biomarker of decreased 11beta-hydroxysteroid dehydrogenase 1 activity in mouse. Br J Pharmacol. 2021;178(16):3309–26.

59. Gomez C, Alimajstorovic Z, Othonos N, Winter DV, White S, Lavery GG, et al. Identification of a human blood biomarker of pharmacological 11beta-hydroxysteroid dehydrogenase 1 inhibition. Br J Pharmacol. 2024;181(5):698–711.

60. Jefcoate CR, McNamara BC, Artemenko I, Yamazaki T. Regulation of cholesterol movement to mitochondrial cytochrome P450scc in steroid hormone synthesis. J Steroid Biochem Mol Biol. 1992;43(8):751–67.

61. Cummings DE. Endocrine mechanisms mediating remission of diabetes after gastric bypass surgery. Int J Obes (Lond). 2009;33 Suppl 1:S33–40.

62. Pham S, Gancel A, Scotte M, Houivet E, Huet E, Lefebvre H, et al. Comparison of the effectiveness of four bariatric surgery procedures in obese patients with type 2 diabetes: a retrospective study. J Obes. 2014;2014:638203.

63. Keidar A, Hershkop KJ, Marko L, Schweiger C, Hecht L, Bartov N, et al. Roux-en-Y gastric bypass vs sleeve gastrectomy for obese patients with type 2 diabetes: a randomised trial. Diabetologia. 2013;56(9):1914–8.

64. Heshmati K, Harris DA, Aliakbarian H, Tavakkoli A, Sheu EG. Comparison of early type 2 diabetes improvement after gastric bypass and sleeve gastrectomy: medication cessation at discharge predicts 1-year outcomes. Surg Obes Relat Dis. 2019;15(12):2025–32.

65. Harris DA, Mina A, Cabarkapa D, Heshmati K, Subramaniam R, Banks AS, et al. Sleeve gastrectomy enhances glucose utilization and remodels adipose tissue independent of weight loss. Am J Physiol Endocrinol Metab. 2020;318(5):E678–E88.

66. Valsamakis G, Anwar A, Tomlinson JW, Shackleton CH, McTernan PG, Chetty R, et al. 11beta-hydroxysteroid dehydrogenase type 1 activity in lean and obese males with type 2 diabetes mellitus. J Clin Endocrinol Metab. 2004;89(9):4755–61.

67. Chapman KE, Coutinho AE, Zhang Z, Kipari T, Savill JS, Seckl JR. Changing glucocorticoid action: 11beta-hydroxysteroid dehydrogenase type 1 in acute and chronic inflammation. J Steroid Biochem Mol Biol. 2013;137:82–92.

68. Subramaniam R, Aliakbarian H, Bhutta HY, Harris DA, Tavakkoli A, Sheu EG. Sleeve Gastrectomy and Roux-en-Y Gastric Bypass Attenuate Pro-inflammatory Small Intestinal Cytokine Signatures. Obes Surg. 2019;29(12):3824–32.

69. Katsogiannos P, Kamble PG, Pereira MJ, Sundbom M, Carlsson PO, Eriksson JW, et al. Changes in Circulating Cytokines and Adipokines After RYGB in Patients with and without Type 2 Diabetes. Obesity (Silver Spring). 2021;29(3):535–42.

70. Cappello C, Zwergal A, Kanclerski S, Haas SC, Kandemir JD, Huber R, et al. C/EBPbeta enhances NF-kappaB-associated signalling by reducing the level of IkappaB-alpha. Cell Signal. 2009;21(12):1918–24.

71. Kossintseva I, Wong S, Johnstone E, Guilbert L, Olson DM, Mitchell BF. Proinflammatory cytokines inhibit human placental 11beta-hydroxysteroid dehydrogenase type 2 activity through Ca2+ and cAMP pathways. Am J Physiol Endocrinol Metab. 2006;290(2):E282–8.

72. Han Y, Meng T, Murray NR, Fields AP, Brasier AR. Interleukin-1-induced nuclear factor-kappaB-IkappaBalpha autoregulatory feedback loop in hepatocytes. A role for protein kinase calpha in post-transcriptional regulation of ikappabalpha resynthesis. J Biol Chem. 1999;274(2):939–47.

73. Esteves CL, Verma M, Rog-Zielinska E, Kelly V, Sai S, Breton A, et al. Pro-inflammatory cytokine induction of 11beta-hydroxysteroid dehydrogenase type 1 in A549 cells requires phosphorylation of C/EBPbeta at Thr235. PLoS One. 2013;8(9):e75874.

74. Kershaw EE, Morton NM, Dhillon H, Ramage L, Seckl JR, Flier JS. Adipocyte-specific glucocorticoid inactivation protects against diet-induced obesity. Diabetes. 2005;54(4):1023–31.

75. Chapman K, Holmes M, Seckl J. 11beta-hydroxysteroid dehydrogenases: intracellular gate-keepers of tissue glucocorticoid action. Physiol Rev. 2013;93(3):1139–206.

76. Scurry MT, Shear L. Stop-flow analysis of the reabsorption of cortisol. Endocrinology. 1969;84(3):681–2.

77. Hall JE, do Carmo JM, da Silva AA, Wang Z, Hall ME. Obesity-induced hypertension: interaction of neurohumoral and renal mechanisms. Circ Res. 2015;116(6):991–1006.

78. Cohen JB. Hypertension in Obesity and the Impact of Weight Loss. Curr Cardiol Rep. 2017;19(10):98.

79. Ebadinejad A, Shahshahani M, Hosseinpanah F, Ghazy F, Khalaj A, Mahdavi M, et al. Comparison of hypertension remission and relapse after sleeve gastrectomy and one-anastomosis gastric bypass: a prospective cohort study. Hypertens Res. 2023;46(5):1287–96.

80. Nudotor RD, Canner JK, Haut ER, Prokopowicz GP, Steele KE. Comparing remission and recurrence of hypertension after bariatric surgery: vertical sleeve gastrectomy versus Roux-en-Y gastric bypass. Surg Obes Relat Dis. 2021;17(2):308–18.

81. Arriza JL, Weinberger C, Cerelli G, Glaser TM, Handelin BL, Housman DE, et al. Cloning of human mineralocorticoid receptor complementary DNA: structural and functional kinship with the glucocorticoid receptor. Science. 1987;237(4812):268-75.

82. Edwards CR, Stewart PM, Burt D, Brett L, McIntyre MA, Sutanto WS, et al. Localisation of 11 beta-hydroxysteroid dehydrogenase--tissue specific protector of the mineralocorticoid receptor. Lancet. 1988;2(8618):986-9.

83. Ferrari P. The role of 11beta-hydroxysteroid dehydrogenase type 2 in human hypertension. Biochim Biophys Acta. 2010;1802(12):1178–87.

84. Funder JW, Pearce PT, Smith R, Smith AI. Mineralocorticoid action: target tissue specificity is enzyme, not receptor, mediated. Science. 1988;242(4878):583-5.

85. Dudenbostel T, Calhoun DA. Use of Aldosterone Antagonists for Treatment of Uncontrolled Resistant Hypertension. Am J Hypertens. 2017;30(2):103–9.

86. Karns AD, Bral JM, Hartman D, Peppard T, Schumacher C. Study of aldosterone synthase inhibition as an add-on therapy in resistant hypertension. J Clin Hypertens (Greenwich). 2013;15(3):186–92.

87. de Souza F, Muxfeldt E, Fiszman R, Salles G. Efficacy of spironolactone therapy in patients with true resistant hypertension. Hypertension. 2010;55(1):147–52.

88. Ueda K, Nishimoto M, Hirohama D, Ayuzawa N, Kawarazaki W, Watanabe A, et al. Renal Dysfunction Induced by Kidney-Specific Gene Deletion of Hsd11b2 as a Primary Cause of Salt-Dependent Hypertension. Hypertension. 2017;70(1):111–8.

89. Uslar T, Newman AJ, Tapia-Castillo A, Carvajal CA, Fardella CE, Allende F, et al. Progressive 11beta-Hydroxysteroid Dehydrogenase Type 2 Insufficiency as Kidney Function Declines. J Clin Endocrinol Metab. 2024.

90. Nogueira EF, Rainey WE. Regulation of aldosterone synthase by activator transcription factor/cAMP response element-binding protein family members. Endocrinology. 2010;151(3):1060–70.

91. Williams LJ, Lyons V, MacLeod I, Rajan V, Darlington GJ, Poli V, et al. C/EBP regulates hepatic transcription of 11beta –hydroxysteroid dehydrogenase type 1. A novel mechanism for cross-talk between the C/EBP and glucocorticoid signaling pathways. J Biol Chem. 2000;275(39):30232–9.

92. Ignatova ID, Kostadinova RM, Goldring CE, Nawrocki AR, Frey FJ, Frey BM. Tumor necrosis factor-alpha upregulates 11beta-hydroxysteroid dehydrogenase type 1 expression by CCAAT/enhancer binding protein-beta in HepG2 cells. Am J Physiol Endocrinol Metab. 2009;296(2):E367–77.

93. Khan S, Livingstone DEW, Zielinska A, Doig CL, Cobice DF, Esteves CL, et al. Contribution of local regeneration of glucocorticoids to tissue steroid pools. J Endocrinol. 2023;258(3).

94. Zhao S, Lin H, Li W, Xu X, Wu Q, Wang Z, et al. Post sleeve gastrectomy-enriched gut commensal Clostridia promotes secondary bile acid increase and weight loss. Gut Microbes. 2025;17(1):2462261.

95. Dzyakanchuk AA, Balazs Z, Nashev LG, Amrein KE, Odermatt A. 11beta-Hydroxysteroid dehydrogenase 1 reductase activity is dependent on a high ratio of NADPH/NADP(+) and is stimulated by extracellular glucose. Mol Cell Endocrinol. 2009;301(1-2):137–41.

96. Wang Y, Nakagawa Y, Liu L, Wang W, Ren X, Anghel A, et al. Tissue-specific dysregulation of hexose-6-phosphate dehydrogenase and glucose-6-phosphate transporter production in db/db mice as a model of type 2 diabetes. Diabetologia. 2011;54(2):440–50.

97. Sandeep TC, Andrew R, Homer NZ, Andrews RC, Smith K, Walker BR. Increased in vivo regeneration of cortisol in adipose tissue in human obesity and effects of the 11beta-hydroxysteroid dehydrogenase type 1 inhibitor carbenoxolone. Diabetes. 2005;54(3):872–9.

98. Finken MJJ, Wirix AJG, von Rosenstiel-Jadoul IA, van der Voorn B, Chinapaw MJM, Hartmann MF, et al. Role of glucocorticoid metabolism in childhood obesity-associated hypertension. Endocr Connect. 2022;11(7).

99. Hua R, Wang GZ, Shen QW, Yang YP, Wang M, Wu M, et al. Sleeve gastrectomy ameliorated high-fat diet (HFD)-induced non-alcoholic fatty liver disease and upregulated the nicotinamide adenine dinucleotide +/ Sirtuin-1 pathway in mice. Asian J Surg. 2021;44(1):213–20.

100. Myronovych A, Kirby M, Ryan KK, Zhang W, Jha P, Setchell KD, et al. Vertical sleeve gastrectomy reduces hepatic steatosis while increasing serum bile acids in a weight-loss-independent manner. Obesity (Silver Spring). 2014;22(2):390–400.

101. Pardina E, Baena-Fustegueras JA, Catalan R, Galard R, Lecube A, Fort JM, et al. Increased expression and activity of hepatic lipase in the liver of morbidly obese adult patients in relation to lipid content. Obes Surg. 2009;19(7):894–904.

102. Gout J, Tirard J, Thevenon C, Riou JP, Begeot M, Naville D. CCAAT/enhancer-binding proteins (C/EBPs) regulate the basal and cAMP-induced transcription of the human 11beta-hydroxysteroid dehydrogenase encoding gene in adipose cells. Biochimie. 2006;88(9):1115–24.

103. Alberti L, Girola A, Gilardini L, Conti A, Cattaldo S, Micheletto G, et al. Type 2 diabetes and metabolic syndrome are associated with increased expression of 11beta-hydroxysteroid dehydrogenase 1 in obese subjects. Int J Obes (Lond). 2007;31(12):1826–31.

104. Wang J, Wang Y, Liu L, Lutfy K, Friedman TC, Liu Y, et al. Lack of adipose-specific hexose-6-phosphate dehydrogenase causes inactivation of adipose glucocorticoids and improves metabolic phenotype in mice. Clin Sci (Lond). 2019;133(21):2189–202.

105. Aragao-Santiago L, Gomez-Sanchez CE, Mulatero P, Spyroglou A, Reincke M, Williams TA. Mouse Models of Primary Aldosteronism: From Physiology to Pathophysiology. Endocrinology. 2017;158(12):4129–38.

